# Relictual Hybridization and Biogeography of Massasauga Rattlesnakes (*Sistrurus* spp.)

**DOI:** 10.1101/2022.08.29.505772

**Authors:** Bradley T. Martin, Marlis R. Douglas, Tyler K. Chafin, John S. Placyk, Stephen P. Mackessy, Jeffrey T. Briggler, Michael E. Douglas

## Abstract

Climate change inevitably leaves behind a genetic footprint within phylogeographic legacies of affected species, as individuals are driven to either disperse to track suitable conditions or adapt *in situ*. One potential consequence is the possibility of hybridization among species, as both geographic ranges and adaptive landscapes shift. The admixture resulting from these newly formed ‘contact zones’ has various outcomes, to include the creation of new lineages. Interpreting these within the context of historic climate change provides clues necessary to predict biotic responses (and thus evolutionary trajectories) as a function of contemporary shifts. Herein, we dissect historic contact zones for Massasaugas (Viperidae; *Sistrurus* spp.) within two distinct North American regions (southwestern United States and Central Great Plains) using ddRAD sequencing. We identified fine-scale but previously unrecognized population structure within the southwestern contact zone, where we detected contemporary intergradation between Prairie and Desert massasaugas (*S. tergeminus tergeminus*, and *S. t. edwardsii,* respectively), with primary divergence indicated by demographic model selection. Within the Central Great Plains, we found evidence for historic secondary contact via Quaternary climatic cycles, subsequently followed by range expansion at the suture zone separating *S. tergeminus* and *S. catenatus*. Extant Missouri populations represent ancestral/relictual vestiges of this earlier hybridization, isolated between the eastern terminus of *S. t. tergeminus* and the western edge of *S. catenatus*. Our results illustrate how abrupt climate change has driven ancestral hybridization, cryptic diversity, and range dynamism within *Sistrurus*.

## 1. Introduction

Species distributions in North America display widespread dynamism in response to historic climatic fluctuations that, in some cases, promoted semi-permeable zones of transition (Antonelli, 2017). Major Quaternary range expansions and contractions, which were punctuated by variable temperature regimes, largely varied in time and place and were modulated by glacial cycles, often with rapid onsets and/or declines of the glacial extent (Clark et al., 1995; Adams et al., 1999; Hewitt, 1996, 1999, 2001; National Research Council, 2002). While some species were substantially displaced by glaciation, followed subsequently by a retreat into and/or emergence from isolated refugia, others adapted *in situ* (Becker et al., 2013). These glacially induced biogeographic processes often occurred iteratively as the climate waxed and waned with ample opportunities for secondary contact during repeated glacial-interglacial cycles. Temperature shifts in these transitions often occurred quite rapidly over geologic time (National Research Council, 2002), with strong and subsequent impacts on biodiversity (Douglas et al., 2009).

Overlapping contemporary ranges (i.e., ‘contact zones’) seemingly reflect spatial transitions between species that promoted intergradation or hybridization (Anderson, 1949; Remington, 1968) and may represent secondary contact via climate-mediated range contractions/expansions (Hewitt, 1996, 1999) or simply an amplified porosity of species-boundaries as driven by transition of habitats (Rhymer and Simberloff, 1996). In any case, such zones are now interspersed throughout North America (Remington, 1968; Swenson and Howard, 2005), but the species-specific context of each varies as a function of local environments and genomic context, as demonstrated in a hybrid zone between Eastern (*Terrapene carolina carolina*) and Gulf Coast (*T. c. major*) North American box turtles. Specifically, directional introgression occurred between *T. c. carolina* and *T. c. major,* whereas hybrid zones composed of either of the *T. carolina* forms and Three-toed box turtle (*T. mexicana triunguis*) displayed strong signatures of selection against interspecific heterozygotes (Martin et al. 2020), reflecting habitat specificity as mediated by the natural history and the genomic context of hybrid zone participants. Linking these interactions with ongoing anthropogenic climatic change is obviously important for pro-active management plans, but baseline data are needed on specific environmental dimensions, underlying adaptive gradients, and the genomic architecture of adaptation. In short, the solution becomes increasingly multifaceted and compounded in lockstep with an enhancement of our evolutionary perspective.

Hybrid zones have long been interpreted as ‘natural laboratories’ (Hewitt, 1988), in that signatures of selection within an ecological gradient can be defined from the extent of introgression expressed relative to a background of genomic homogenization (Payseur, 2010; Taylor and Larson, 2019). This framework can be extended by contrasting taxon-pairs within a speciation continuum, or occupying different environmental niches, thus offering a ‘window’ from which to view the numerous extrinsic factors such as climate change that drive species-interactions (e.g., Taylor et al., 2015). Essentially, it is a necessary first step in disentangling the quagmire of context-specific factors that govern individual outcomes.

Cataloguing biotic responses to climatic change and subsequently assaying for genomic adaptations are now increasingly common approaches, due largely to reduced sequencing costs and an ever-increasing genomic ‘database’ for non-model organisms. Importantly, such resources offer a predictive capacity for hypothesis-testing going forward, particularly regarding the unrelenting changes characterizing the Anthropocene. We follow this trajectory herein by quantifying the formation of contact zones as they relate to the historic biogeography of viperid snakes (*Sistrurus* spp.), whose distributions span much of the southwestern, midwestern, and eastern United States and southeastern Canada.

### 1.1. Study species

The genus *Sistrurus* (Garman 1883) contains the Massasauga and Pygmy rattlesnakes (Viperidae: Crotalinae) and forms a sister clade to the remaining North American rattlesnakes (*Crotalus*; Murphy et al., 2002; Castoe and Parkinson, 2006). *Sistrurus* inhabits much of the eastern United States, the Great Plains from Iowa to southernmost Texas, and the shortgrass prairie in central Texas, New Mexico, and southeastern Arizona (Fig. 1). It contains three currently recognized species: Eastern (*S. catenatus*) and Western *(S. tergeminus*) massasaugas, and the Pygmy Rattlesnake (*S. miliarius*). *Sistrurus catenatus* is monotypic, whereas *S. miliarius* and *S. tergeminus* include subspecies: The Carolina (*S. m. miliarius*), Dusky (*S. m. barbouri*), and Western Pygmy (*S. m. streckeri*) Rattlesnakes, and the Prairie (*S. t. tergeminus*) and Desert (*S. t. edwardsii*) Massasaugas. The latter two contain highly disjunct populations in Missouri, Colorado, New Mexico, and southeastern Arizona that may correspond to ESUs (evolutionarily significant units) and MUs (management units) (Greene, 1997), although there is disagreement about these designations (Bylsma et al., 2021). Their conservation status has been mediated by anthropogenically-induced habitat loss and fragmentation, road-based mortality, and prescribed fires (Szymanski, 1998; Szymanski et al., 2016), particularly for *S. catenatus* and *S. tergeminus* ssp., although a fragmented habitat preceding anthropogenic impacts is also acknowledged (Chiucchi and Gibbs, 2010; Sovic et al., 2019; Ochoa et al., 2020). Herein our focus is on *S. catenatus* and *S. tergeminus*.

**Figure 1:**
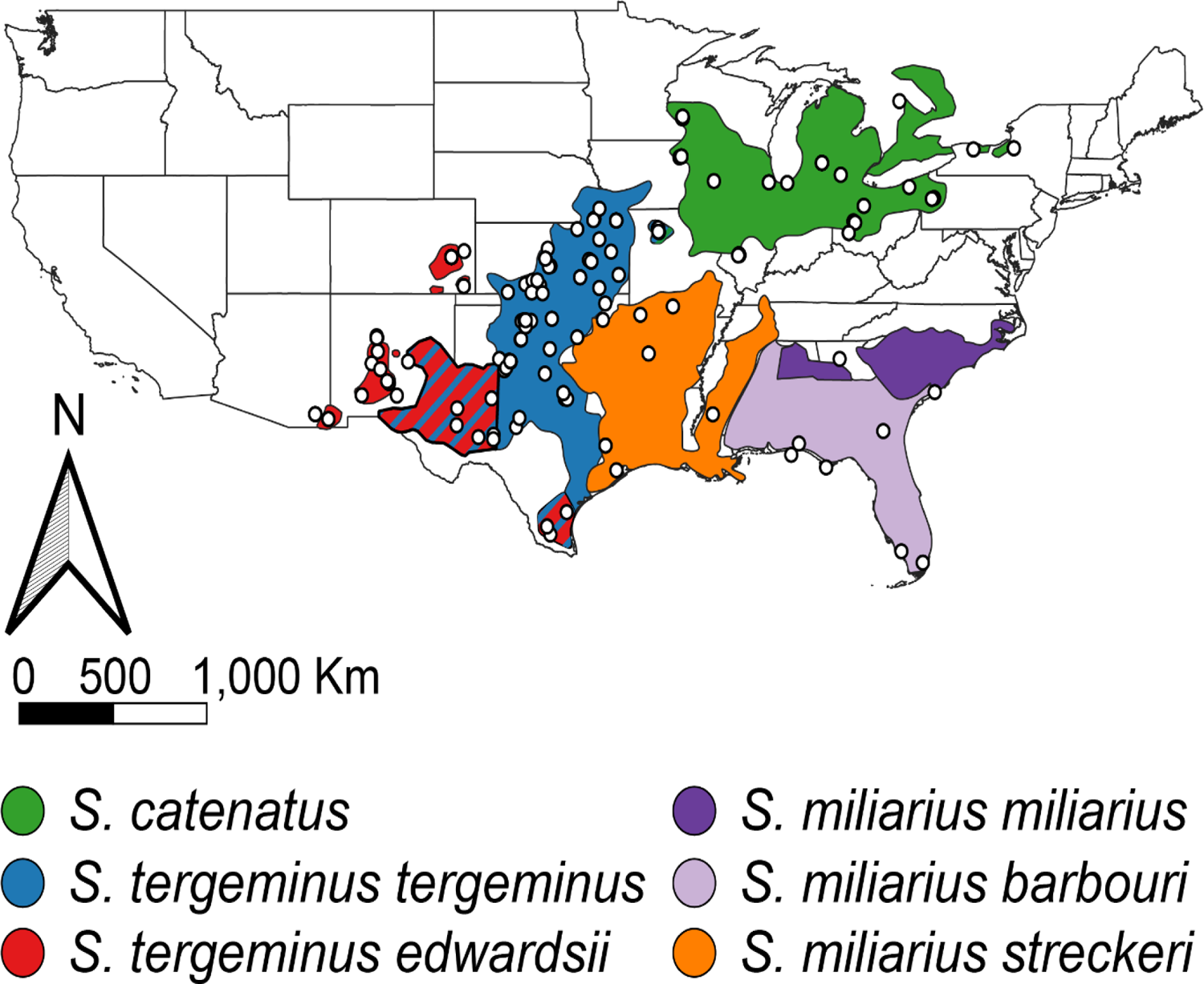
*Sistrurus* range map depicting subspecies distributions and ddRAD sequencing samples. Cross-hatched areas in Texas, New Mexico, and Missouri indicate overlapping ranges with known contact zones. White circles indicate samples sequenced for this study. Adapted from Sanz et al. (2006).

Recent phylogenetic assessments (e.g., Kubatko et al., 2011; Hein, 2015) found *S. catenatus* monophyletic and *S. t. tergeminus* paraphyletic with respect to *S. t. edwardsii*. Population structure is weak within *S. catenatus* (Chiucchi and Gibbs, 2010; Sovic et al., 2016), whereas assessments within *S. tergeminus* have been inconsistent. Here, Ryberg et al. (2015) only supported the differentiation of *S. t. tergeminus* in Missouri and Oklahoma, whereas in *S. t. edwardsii,* non-admixed populations were identified in New Mexico between the Pecos River and Arizona and at the southern tip of Texas. Both subspecies lacked population structure in Oklahoma/ western Texas/eastern New Mexico and Colorado/ Kansas. Likewise, Anderson et al. (2009) observed distinct populations of *S. t. edwardsii* in Arizona and New Mexico, whereas Bylsma et al. (2021) suggested *S. tergeminus* represented but a single gene pool without subspecific diversification. Nevertheless, the Missouri and western Texas/eastern New Mexico populations may indeed overlie contact zones (Minton, 1983), each with a separate chronology.

Hybridization in Missouri the population of *Sistrurus* has likewise been suggested, but with evaluations again lacking consistency. Individuals from transitional habitat were morphologically-identified as intermediate between *S. catenatus* and *S. t. tergeminus* (Evans and Gloyd, 1948), although recent genetic studies have designated them as pure *S. t. tergeminus* (Gerard et al., 2011; Gibbs et al., 2011). However, a neighboring, disjunct Iowa population did display signatures of introgression between the two subspecies (Sovic et al., 2016). Given the contemporary nature of the Iowa population, its observed levels of introgression are likely historic. The Missouri population may reflect a similar situation, but this would require greater genomic resolution than previously afforded (e.g., Chafin et al., 2019).

There is also anecdotal evidence for potential admixture between *S. tergeminus* subspecies, particularly given the presence of intermediate habitat in the contact zone separating western Texas/eastern New Mexico. Few studies have assessed genetic population structure in *S. tergeminus* ssp., and of these, most employed single-gene markers and/or microsatellite (msat) DNA. The utilization of next generation sequencing (NGS) would provide a more fine-scaled population structure for these clades and allow a much more focused interpretation of their phylogeographies and potential admixtures. We do so herein, by employing large samples drawn across a broad geographic range, from which we derive a dataset comprised of single nucleotide polymorphisms (SNPs). This allowed us to explore the demographic history of *Sistrurus* within its recognized contact zones, and to draw inferences regarding the climatic and biogeographic processes that have shaped its evolutionary history.

## 2. Materials and methods

### 2.1. Study system, tissue collection, and DNA extraction

Broadly distributed samples (*N*=226) were acquired (Fig. 1) by targeting each recognized species and subspecies across each U.S. state within which they occur. We particularly focused on disjunct Missouri populations, as well as the contact zone in the southwestern United States. Our sampling procedures were approved under Institutional Animal Care & Use Committee (Arizona State University: 93–280R). DNA was extracted either from shed skins or non-invasive blood samples (Supplementary Table S1) using PureGene® or DNeasy Blood and Tissue Kits (Qiagen), and genomic DNA stored at −20LC. DNA quantification was performed using broad-range DNA fluorometry (Qubit; Thermo Fisher Scientific), with each extract verified for sufficiently high molecular weight DNA via 2% agarose gel electrophoresis.

### 2.2. Library preparation and bioinformatics

Genomic DNA (∼500-1,000ng) was digested at 37LC using *PstI* (5’-CTGCA|G-3’) and *MspI* (5’-C|CGG-3’). Digestion was verified via 2% agarose gel electrophoresis, then purified (1.5X AMPure XP magnetic beads; Beckman Coulter). DNA was standardized to 100ng, ligated with uniquely barcoded adapters, and pooled into 48-individual libraries. Following ligation, we size-selected 323-423bp using a Pippin Prep (Sage Science), followed by a 12-cycle polymerase chain reaction (PCR) using high-fidelity Phusion DNA Polymerase (New England BioLabs). DNA amplification was confirmed using Qubit fluorometry, and quality control steps (qPCR and fragment visualization) were performed at the Genomics and Cell Characterization Core Facility (University of Oregon/Eugene). Samples were sequenced in pools distributed across two single-end 100bp lanes on the Illumina Hi-Seq 4000.

Raw sequence quality was first assessed (FASTQC v0.11.5; Andrews, 2010), with reads then demultiplexed, clustered, and aligned (IPYRAD v0.7.30; Eaton and Overcast, 2020). We removed barcodes and adapters, then assembled our alignments *de novo*, with a minimum coverage depth requirement of ≥20 reads per fragment and a missing data threshold of 50%. Sites with >75% heterozygosity were removed as potential paralogs, and the last five bases trimmed from each locus to mitigate potential sequencing error due to primer degradation. Various clustering thresholds were tested to determine the optimum, ranging from 70% to 97% [CLUSTOPT (McCartney-Melstad et al., 2019) via R v3.6.3 (R Development Core Team, 2018)]. These were evaluated across three metrics: 1) Cumulative variance from a principal component analysis (PCA); 2) Pearson’s correlation coefficients between genetic distance versus percent missing data; and 3) isolation-by-distance (IBD). The optimal threshold was interpreted as the inflection point in the resulting slopes. Finally, we applied a post-hoc alignment filtering via a custom script (*nremover.pl*; https://github.com/tkchafin/scripts) to remove poorly sequenced individuals (i.e., >90% missing data).

### 2.3. Phylogenomic analysis of Sistrurus

We first inferred a maximum-likelihood (ML) tree of concatenated loci (IQ-TREE v2.0.6; Minh et al., 2020) using the edge-linked partition model (Chernomor et al., 2016), with members of *Crotalus* and *Agkistrodon* as outgroups (Parkinson et al., 2002). Loci lacking parsimony-informative sites were removed, with the remaining full loci being concatenated into a single NEXUS file and demarcated using a per-locus partition block, as executed with three custom scripts [*filterLoci.py*, *concatenateNexus.py*, and *filterUninformative.py* (https://github.com/tkchafin/scripts and https://github.com/btmartin721/ddrad_scripts)].

Substitution models were assessed on a per-partition basis. Those spanning equivalent models (=individual loci) were merged using the fast-relaxed clustering method (‘rclusterf’ option). From the top 10% of clusters, we then obtained the optimal nucleotide substitution model for each selected super-partition (MODELFINDER; Kalyaanamoorthy et al., 2017). Tree support was assessed with 1,000 ultrafast bootstrap replicates (UFBOOT; Hoang et al. 2017), with the ‘bnni’ and ‘safe’ options to mitigate overestimation of support and prevent numerical underflow, respectively. This topology was further contrasted with phylogenetic networks inferred using the SNaQ method in PHYLONETWORKS (see below).

### 2.4. Divergence dating

As a baseline for assessing the demographic history of *Sistrurus*, we estimated divergence times from the IQ-TREE phylogeny using the LSD2 approach (least square dating v2, IQ-TREE v2.1.2; To et al., 2016). Molecular clock calibrations included fossil-based minimum/maximum age constraints for four nodes, with lower bounds supplied for the MRCA (most recent common ancestor). These are: *Sistrurus* [(9.0 Mya) (Parmley and Holman, 2007)]; *Agkistrodon contortrix* + *A. piscivorous* [(4.5 Mya) (Holman, 2000; Guiher and Burbrink, 2008; Douglas et al., 2009)]; *Crotalus* + *Sistrurus* [(15.5 Mya) (Parmley and Holman, 2007)]; and *Agkistrodon* + *Crotalus* [(7.0 Mya) (Douglas et al., 2006, 2009)]. A maximum root constraint of 22.0 Mya represented the earliest known pit viper fossil (Holman, 2000; Parmley and Holman, 2007, Douglas et al., 2009). To calculate confidence intervals for divergence times, LSD2 was run on 1,000 bootstrapped trees with branch lengths simulated from a relaxed-clock Poisson distribution. The resulting phylogeny was then plotted using R packages [PHYTOOLS (Revell, 2012); GGTREE (Yu et al., 2017)].

### 2.5. Phylogeographic signal within Sistrurus

Based on visual inspection of phylogenetic patterns, we explored our time-calibrated phylogenetic signal using a Brownian motion evolutionary model corresponding to the observed spatial signal (PICANTE R-package; Kembel et al., 2010). Phylogenetic-spatial signal was tested using the *multiPhylosignal* function for all combinations of adjacent *Massasauga* taxa (i.e., *S. catenatus*, *S. t. tergeminus*, and *S. t. edwardsii*, plus the whole tree). Signal was significantly influenced by latitude and longitude when *K*-statistic (Blomberg et al., 2003) was >1.0 and the phylogenetic signal non-random and greater than expected (*P* < 0.05).

We also estimated the distance of each tip of the phylogeny from the most ancestral node per subspecies (PALEOTREE R-package; Bapst, 2012). Here, the time-calibrated IQ-TREE phylogram was input into the ‘dateNodes’ PALEOTREE function, and the number of tips calculated between the current one and the clade root. Nodal distances were standardized as proportions to account for differences in clade depth.

### 2.6. Admixture and population structure

We assessed population structure and admixture [ADMIXTURE v1.3.0 (Alexander and Lange, 2011) via ADMIXPIPE (Mussmann et al., 2020b)] to identify contact zones and potentially pure parental populations that may correspond to ESUs or MUs. Sites either monomorphic or with a minor allele frequency <1% were *a priori* removed from the unlinked IPYRAD SNP output (Linck and Battey, 2019), using filtering options in ADMIXPIPE. We ran ADMIXTURE for *K*=1-10 clusters with 20 replicates per *K* and 20-fold cross-validation (CV). Optimal *K* was chosen as having the lowest CV score, and the ADMIXTURE results were visualized as stacked bar-plots (ADMIXPIPE scripts *cvSum.py* and *distructRerun.py*). ADMIXTURE proportions were plotted as pie charts on a range map of *Sistrurus,* allowing the spatial distributions of derived clusters to be visualized (QGIS v3.16; QGIS Development Team, 2020).

### 2.7. Tests of hybridization and deep-time reticulation

We employed four-taxon *D*-statistic tests to infer if our observed admixture was statistically supported [Green et al. 2010; Durand et al. 2011; via COMP-D (Mussmann et al., 2020a)]. As with ADMIXTURE, singletons and monomorphic sites were excluded. We employed a custom script to generate the inputs for COMP-D (*makeCompD.py*; https://github.com/tkchafin/makeCompD). The parallel MPI (message passing interface) version of COMP-D was run with 1,000 bootstrap replicates and heterozygous sites included (‘hinclude’ option). Benjamini and Yekutieli (B-Y) corrections were independently applied to adjust *P*-values for multiple comparisons (Bonferroni, 1936; Benjamini and Yekutieli, 2001). Positive *D*-statistic tests were then summarized per individual and population.

We also inferred phylogenetic networks and computed ancestry components to refine hypothesized reticulations previously identified with ADMIXTURE and the *D*-statistic. We did so by using the SNAQ method (PHYLONETWORKS; Solís-Lemus and Ané 2016; Solís-Lemus et al. 2017). To assess support, locus-wise concordance factors (CFs) were calculated using a Bayesian concordance analysis (BUCKy; Larget et al., 2010), run in parallel across quartets (TICR pipeline; Stenz et al., 2015). Given the computational overhead imposed by network inference, we computed our CFs across a reduced dataset of 3,988 loci, chosen as a non-monomorphic subset containing at least five parsimoniously informative sites, and for which at least a single diploid genotype could be sampled per targeted tip. To maximize the number of loci employed, we reduced taxa to eight sampled tips, including all major taxa and intraspecific clades identified in earlier analyses.

Posterior distributions were generated for each gene tree (MRBAYES v3.2.6; Ronquist et al., 2012), with 4 independent chains each and 100,000,000 iterations, 50% of which were discarded as burn-in, and with sampling every 10,000 generations to reduce autocorrelation among samples. Quartet concordance factors (CFs) were then run across all possible four-taxon combinations, using a chain length of 10,000,000 and 50% burn-in. As input to PHYLONETWORKS, a starting tree was first generated (QUARTETMAXCUT; Snir and Rao, 2012), followed by network estimation under models of 0-6 hybrid nodes (*h*). Models were evaluated using 100 independent replicates each, with the best-fit model maximizing first-order change in pseudo-likelihood (ΔL).

Finally, to corroborate our PHYLONETWORKS analysis, we also estimated admixture edges (*m*) along a maximum-likelihood phylogeny [TREEMIX v1.1; Pickrell and Pritchard 2012 via the IPYRAD analysis toolkit (Eaton and Overcast, 2020)]. The input SNP alignment was subsampled to one random non-monomorphic SNP per ddRAD locus, replicated nine times. Optimal *m* was determined for each subsample replicate by observing the inflection point of the log-likelihood scores, using 1,000 bootstrap replicates and the more thorough ‘global’ search option.

### 2.8. Demographic modeling using GADMA

Finally, we assessed introgression events in *S. tergeminus ssp.* by employing demographic modeling [(moments; Jouganous et al., 2017) via the GADMA v1 pipeline (Noskova et al., 2020)]. To estimate the size (N_E_), divergence time (τ), and migration (m) parameters of each population, we assessed a joint site-frequency spectrum (jSFS) to each of our user-defined demographic models. A MOMENTS model search was automated with a genetic algorithm in GADMA to select the optimal model, per the Akaike Information Criterion (AIC). Model search was terminated when log-likelihood values failed to improve over 100 algorithmic iterations, followed by local optimization searches to further refine parameters.

Two GADMA runs were conducted: First, the *Sistrurus* alignment was subset to three populations (the maximum allowed), which included *S. t. tergeminus* from Missouri (=TEMO), *S. t. tergeminus* from all other localities (=TERG), and *S. catenatus* (=CAT). Second, GADMA was run with *S. t. tergeminus* and *S. t. edwardsii*. Sites in each input alignment were filtered to a minimum of 50% coverage in any given population and thinned to one random bi-allelic SNP per ddRAD locus. This random selection was then repeated 100 times to provide non-parametric bootstrap replicates for estimating GADMA parameter confidence intervals (*easySFS.py* script; https://github.com/isaacovercast/easySFS). Alleles in each resulting SFS were down-projected (*easySFS.py*) to yield counts that maximized the number of segregating sites (per the MOMENTS manual). The GADMA model search explored up to two divergence times prior to and following each population split. GADMA requires the population size of the reference population (N_ref_) as a means of scaling the moments parameters in actual time (years), rather than genetic units. This was calculated as: N_ref = θ/θ_0 = (4 × Ne × u × L)/(4 × µ × L), where N_e_ is the effective population size, µ is the mutation rate per generation, and *L* the effective sequence length after filtering, calculated as: L = (total sequence analyzed × SNPs retained in M O M E N T S)/(total SNPs in analyzed sequence).

To calculate a mutation rate [per Gutenkunst et al. (2009) and the GADMA manual], we first derived the following: Average sequence divergence (*D_xy_*) from the input alignment (DNASP v6.12.03; Rozas et al., 2017), a *Sistrurus* generation time (G) of ∼4.0 years (Sovic et al., 2016), and a divergence time for *S. catenatus* x *S. t. tergeminus*, as estimated from the time-calibrated phylogeny. We then calculated the mutation rate as: μ = (Dxy X G)/(2 X r).

## 3. Results

### 3.1. Data assembly and filtering

The IPYRAD clustering threshold was set at 0.82 to maintain a minimal correlation (r ≤ 0.3) between genetic distance and percent missing data in the alignment (Supplementary Fig. S1A). IBD and the cumulative variance from PCA were largely unaffected by clustering thresholds (Supplementary Figs. S1B, S1C). The final IPYRAD alignment included 49,879 parsimoniously informative sites across 10,190 ddRAD loci from N=226 *Sistrurus* (Fig. 1; Supplementary Table S2).

### 3.2. Phylogeographic and demographic analyses

The ML phylogeny indicated reciprocal monophyly for *S. miliarius*, *S. catenatus*, and *S. tergiminus*. Within *S. tergiminus*, we found *S. t. tergeminus* to be paraphyletic with respect to *S. t. edwardsii* (Fig. 2, Supplementary Fig. S2). The *S. tergeminus* clade was largely pectinate, with divergences extending from northeast to southwest. Those most basally divergent were in Missouri, whereas the most derived were in Texas/New Mexico. This longitudinal signal was statistically corroborated (PICANTE; *P*=0.001), with the *K*-statistic >1.0 for all but the whole-tree analysis. The longitudinal signal in the analysis was consistently higher than latitudinal (Table 1), except when *S. t. edwardsii* was independently considered. Finally, PALEOTREE indicated disparate directional patterns of increasingly derived lineages (Fig. 3A), with *S. catenatus* extending northeast and *S. tergiminus* southwest. Individuals with the least nodal distance to the root node in *S. t. tergeminus* and *S. t. edwardsii* were in Kansas/Missouri/Iowa, and Texas/New Mexico, respectively.

**Figure 2:**
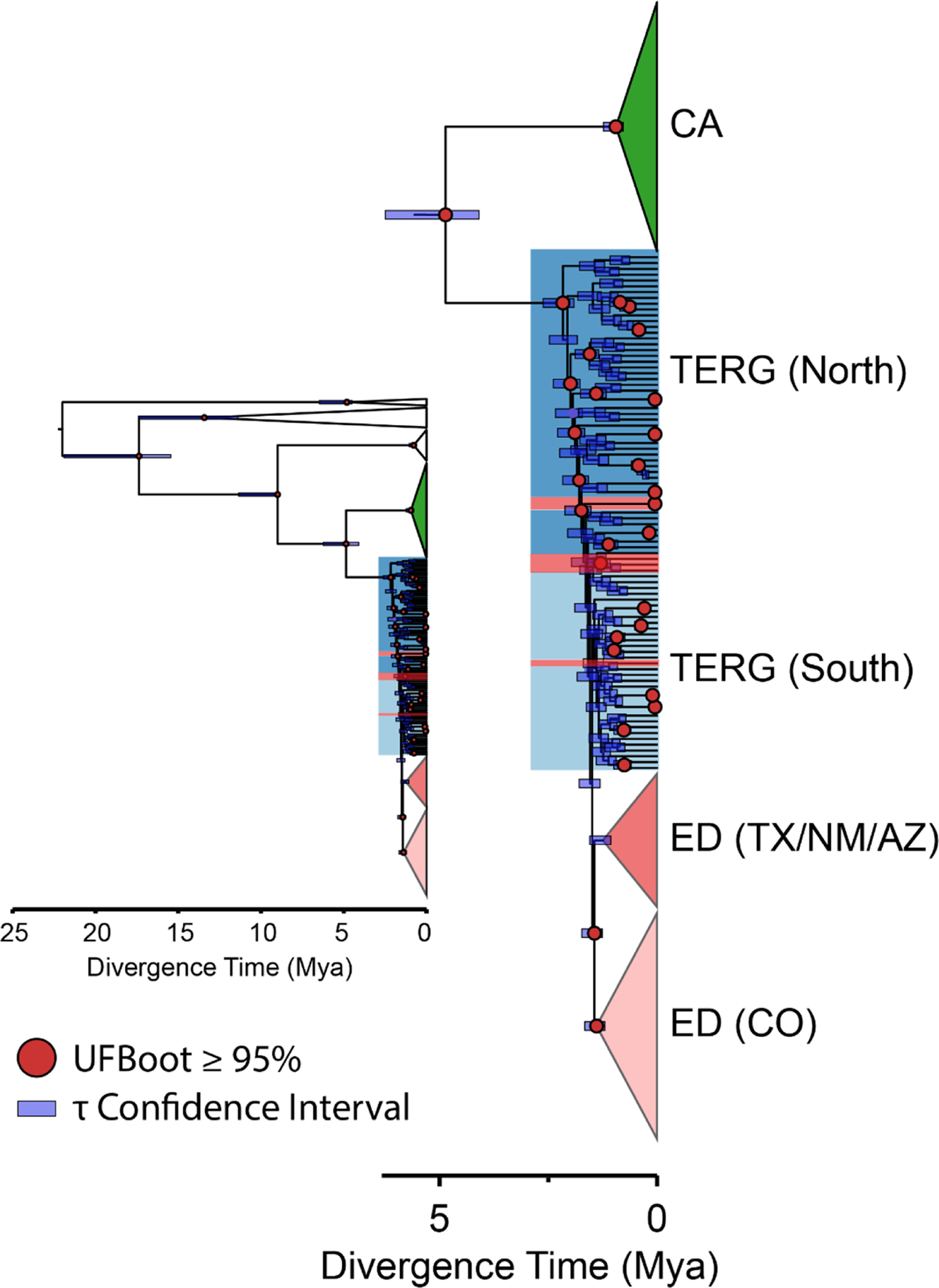
*Sistrurus* molecular clock, with phylogeny inferred using the edge-linked partition model from IQ-TREE v2.1.2 and scaled to actual time (millions of years ago, Mya) using the IQ-TREE implementation of least square dating (LSD). Populations are color-coded according to ADMIXTURE analysis (Fig. 3B), with *S. catenatus* (CA) from all available localities, *S. tergeminus tergeminus* (TERG) subdivided into northern and southern clusters, and *S. t. edwardsii* (ED) represented by Texas/ New Mexico/ Arizona and Colorado populations. Red circles indicate nodes with ultrafast bootstrap (UFBoot) support ≥ 95%, and node bars correspond to divergence time (τ) confidence intervals as calculated from 1,000 input trees with branch lengths simulated from a relaxed-clock Poisson distribution. The main plot is zoomed to *S. catenatus* and *S. tergeminus* ssp., whereas the inset plot shows the full tree including *S. miliarius* and the outgroups (several species of *Agkistrodon* and *Crotalus*) (unfilled triangles).

**Figure 3:**
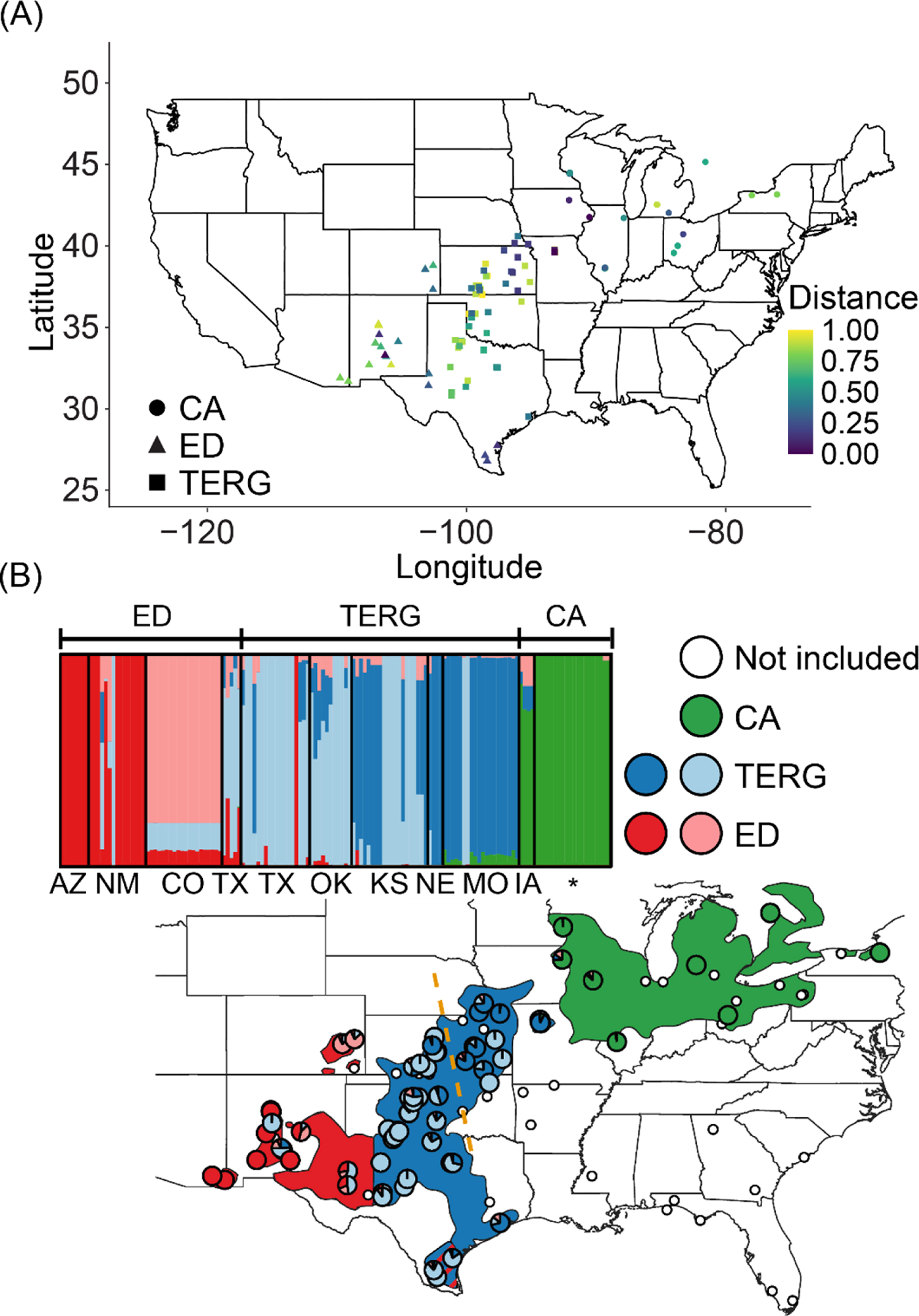
PALEOTREE and ADMIXTURE results for *Sistrurus*. CA=*S. catenatus,* ED=*S. tergeminus edwardsii*, TERG=*S. tergeminus tergeminus*. (A) PALEOTREE diversification distances, with nodal distances = proportion of nodes between tip and most ancestral species node. (B) Pie charts on the map geographically display the ADMIXTURE proportions from the above barplot for *K*=5, with each bar representing one individual and bars with mixed colors indicating mixed ancestry. The orange dashed line illustrates two observed *S. t. tergeminus* populations. The barplot is partitioned into sub-populations based on subspecies field identification (upper guide) and U.S. state locality (bottom guide; IA=Iowa, MO=Missouri, NE=Nebraska, KS=Kansas, OK=Oklahoma, TX=Texas, CO=Colorado, NM=New Mexico, AZ=Arizona, =multiple state localities).

**Table 1:**
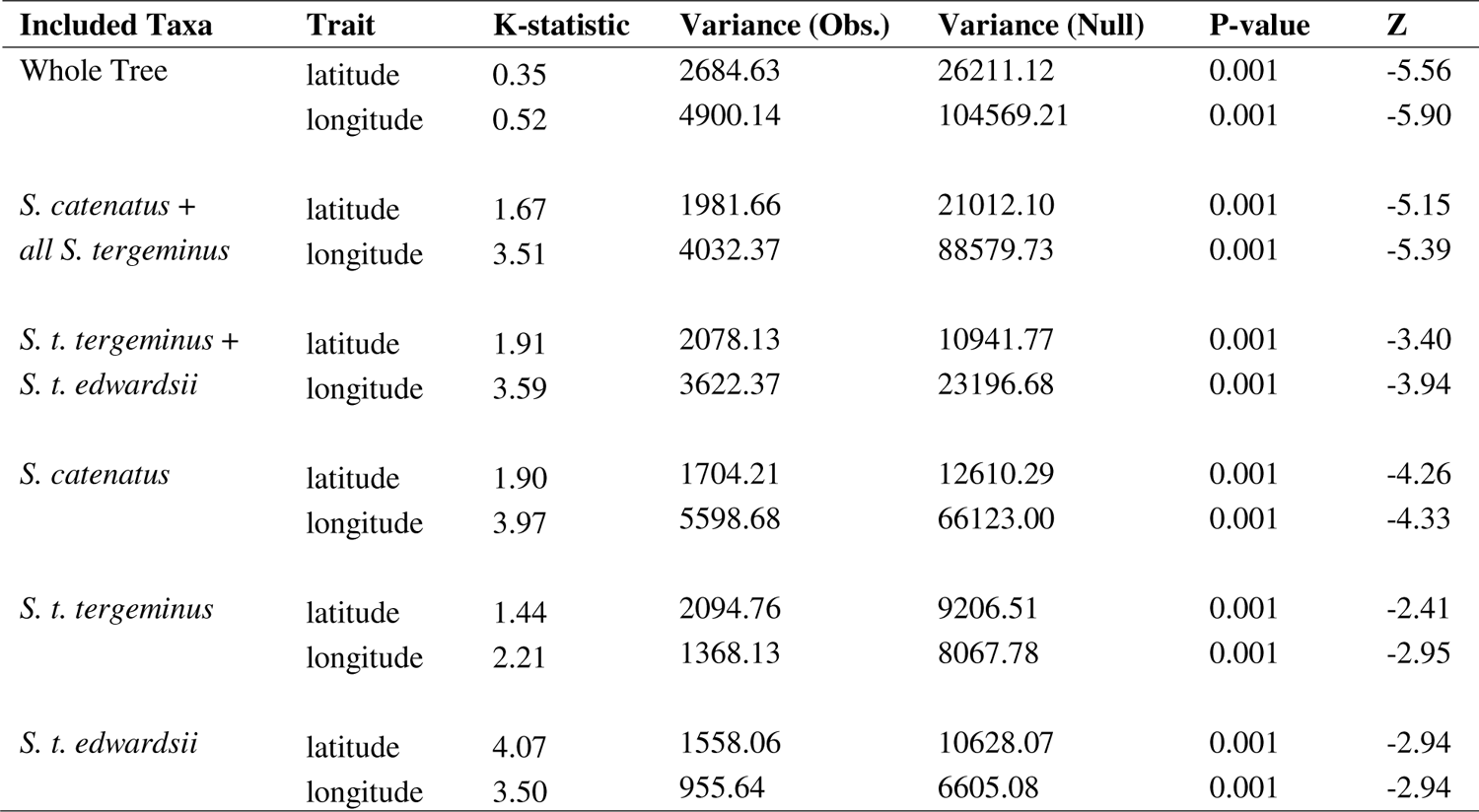
PICANTE analysis assessing spatial signals along a time-calibrated phylogeny (Fig. 3).

The Midwestern GADMA SFS was down-projected to 30 x 30 x 28 alleles for *S. catenatus*, *S. t. tergeminus* (Missouri), and all other *S. t. tergeminus* (hereafter referred to as the ‘CAT x TERG x TEMO’ model). The optimal demographic model for this arrangement included a post-divergence period of weak migration followed by extensive and recent migration (i.e., within the last 7 Kya) (Fig. 4A, Supplementary Fig. S3A). Every supported migration edge and population bottleneck involved *S. catenatus* and *S. t. tergeminus* (Missouri).

**Figure 4:**
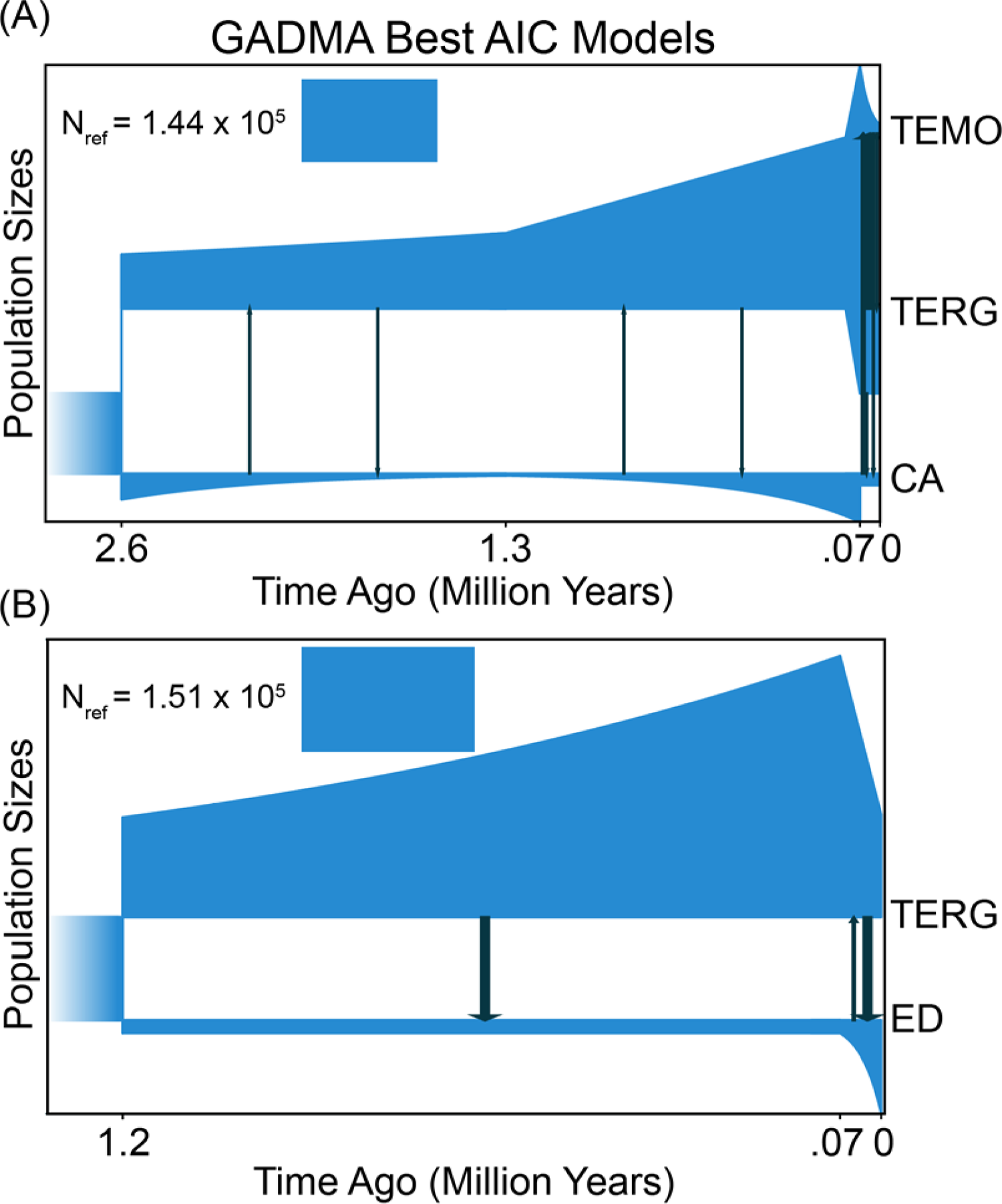
Demographic models chosen by GADMA. Arrows depict migration events with sizes proportional to estimated migration parameters. Branch size on the Y-axis is scaled to the reference population size (N_ref_). (A) Optimal model for *Sistrurus catenatus* (CA) X *S. tergeminus tergeminus* from Missouri (TEMO) and from all other sample localities (TERG). (B) Optimal model for *S. t. tergeminus* from all sample localities except Missouri (TERG) and *S. t. edwardsii* (ED).

Southwestern *S. t. edwardsii*/*S. t. tergeminus* (‘ED x TERG’ model) were down projected to 40 x 56 alleles. In contrast to the CAT x TERG x TEMO model, we noted a strong continuous migration-since-divergence, as well as a recent (<7 Kya) expansion of the *S. t. edwardsii* population (Fig. 4B, Supplementary Fig. S3B).

### 3.3. Hybridization in two North American contact zones

The CV scores from ADMIXTURE (Supplementary Fig. S4) suggested an optimal *K*=5, which included *S. catenatus* and two populations each for *S. t. tergeminus* and *S. t. edwardsii* (Fig. 3B). The two *S. t. tergeminus* populations were generally (albeit porously) localized along a geographic gradient extending from northwest (darker blue shade primarily in northcentral MO/ eastern Kansas/ southeastern Nebraska) to southwest (lighter blue in Kansas/Texas/Oklahoma/New Mexico) (Fig. 3B). The darker blue northwestern population also aligns with the point-of-origin in the PALEOTREE analysis (Fig. 3A).

Similarly, *S. t. edwardsii* was partitioned into populations from Texas/New Mexico/Arizona (darker red) and Colorado (lighter red) (Fig. 3B). Admixture across species/subspecies was localized by region in the midwestern and southwestern contact zones. Results were similar for the *K*=4 admixture analysis (Supplementary Fig. S5), with the exception that *S. t. edwardsii* (CO) instead displayed ∼50% co-ancestry with southern *S. t. tergeminus* and *S. t. edwardsii*.

The four-taxon *D*-statistic tests only supported introgression between *S. t. tergeminus* (MO) and *S. catenatus* (Figs. 5, 6), with significance tests being negligible among southwestern *S. t. tergeminus* and *S. t. edwardsii* (Supplementary Fig. S6). Results from PHYLONETWORKS and TREEMIX concurred (Fig. 6), with a single admixture edge (Supplementary Fig. S7) between *S. t. tergeminus* (MO) + *S. catenatus.* The ancestry proportions (α) from PHYLONETWORKS assigned *S. t. tergeminus* (MO) as 33.8% *S. catenatus*. TREEMIX results differed slightly by only connecting *S. catenatus* (IA) to *S. t. tergeminus* (MO), whereas *D*-statistics and PHYLONETWORKS did so with all *S. catenatus* populations other than IA.

**Figure 5:**
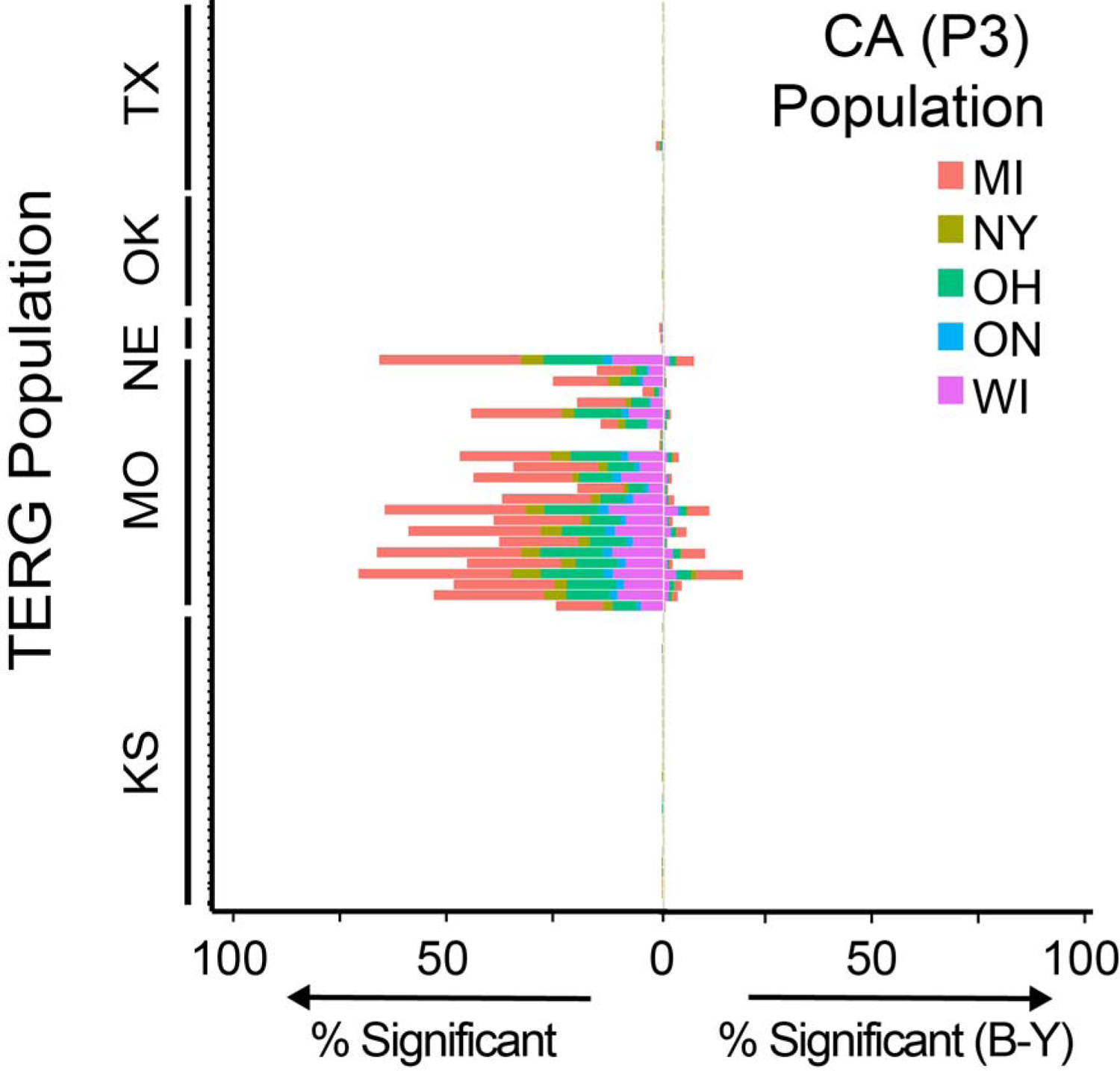
Four-taxon *D*-statistic tests for CA=*Sistrurus catenatus* (P3) and TERG=*S. tergeminus tergeminus* (P1 and P2). Stacked bars indicate the percentage of significant *D*-statistic tests per P3 sample locality with (left) no *P*-value correction for multiple tests and (right) a Benjamini– Yekutieli (B-Y) correction for controlling the false discovery rate. State locality abbreviations include: TX=Texas, OK=Oklahoma, NE=Nebraska, MO=Missouri, KS=Kansas, MI=Michigan, NY=New York, OH=Ohio, WI=Wisconsin, and ON=Ontario (Canada). The outgroups (P4) included *S. miliarius streckeri* and *S. m. barbouri*.

**Figure 6:**
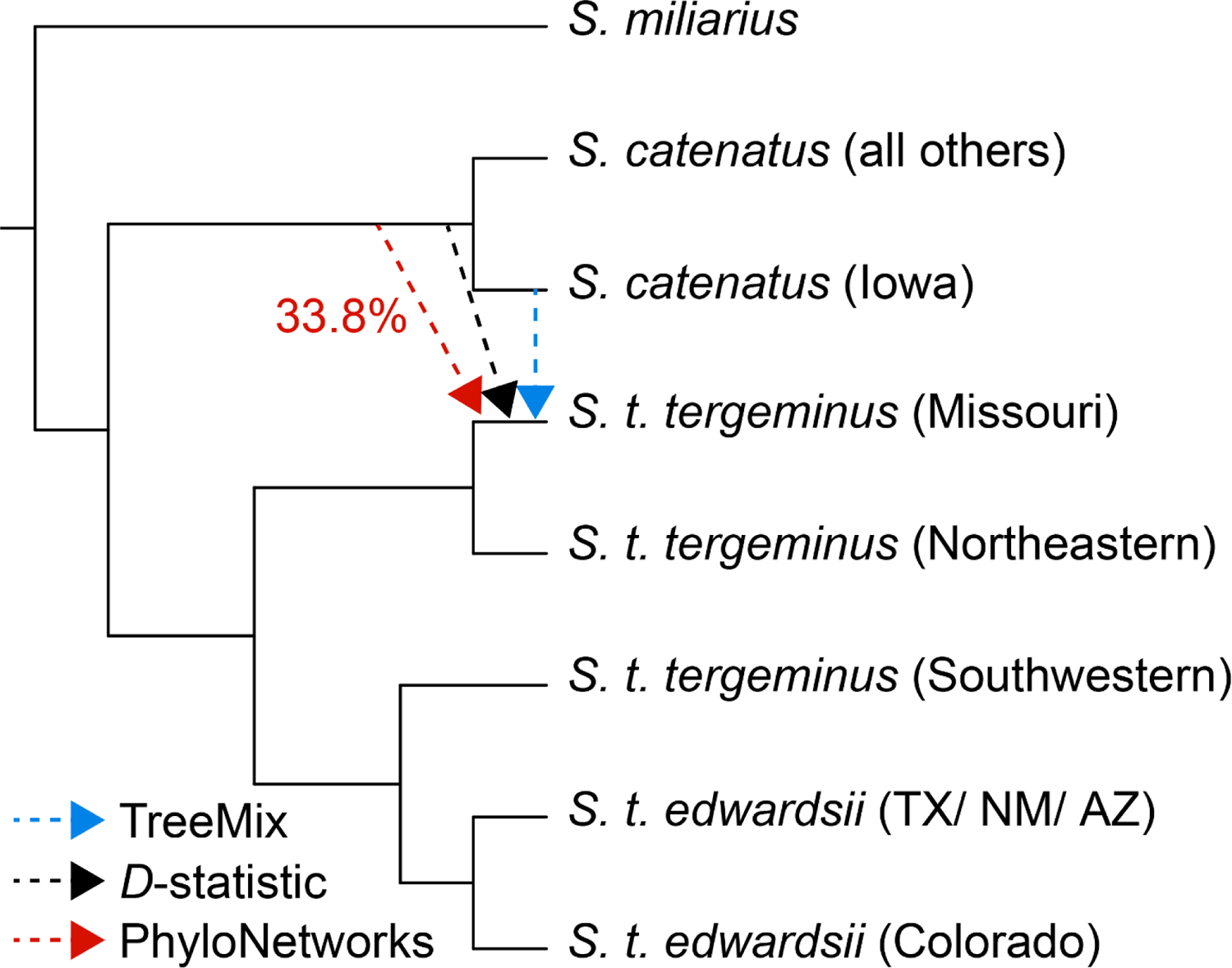
Supported *Sistrurus* admixture edges among TREEMIX, four-taxon *D*-statistic tests, and PHYLONETWORKS. The percentage corresponding to the PHYLONETWORKS (red arrow) illustrates *S. catenatus* ancestry estimated for Missouri *S. t. tergeminus*. Scientific names are followed by state or regional locality. Northwestern and southwestern *S. t. tergeminus* correspond to sub-populations identified in the ADMIXTURE analysis (Fig. 3). TX=Texas, NM=New Mexico, AZ=Arizona.

## 4. Discussion

Our results expanded upon previous studies by employing a greater geographic and genomic breadth of sampling. This allowed us to obtain a more refined picture of both intraspecific population structure and reticulation in *Sistrurus*. Our dissection of the two contact zones underscored the presence of divergent signals within each, with contrasting environmental, physiographic, and temporal processes in play. Hybridization during periods of rapid environmental change provides adaptive variation particularly at the advancing edges of dispersal. Here, local adaptation will hinge upon population demography as well as genetic architecture (Jones et al., 2020). We discuss the demographic patterns inherent within the environmental, physiographic, and temporal processes of our study system, and identify evolutionary and conservation implications that have subsequently emerged.

### 4.1. Historical introgression between S. catenatus and S. tergeminus

In Missouri, 22 (of 24) *S. t. tergeminus* (92%) from the three remaining inhabited counties (Linn, Holt, and Chariton) were identified as hybrids, with the network analysis attributing 33.8% ancestry to *S. catenatus*. Our results concur with a previous morphological assessment establishing these populations as intermediate (Evans and Gloyd, 1948). Our results also stand in contrast with simulation models employed in more recent molecular studies (Gerard et al., 2011; Gibbs et al., 2011). Given that Missouri populations are disjunct from larger (and contiguous) populations of *S. catenatus* and *S. t. tergeminus*, potential introgression is inferred to be historic.

In this study, genetic signatures of historic admixture were clearly detected in the Missouri populations, due largely to the greater capacity of SNP data to diagnose evolutionary and demographic processes at deeper time scales (Morin et al., 2004; Haasl and Payseur, 2011; Lee et al., 2018; Chafin et al., 2021). In addition, numerous studies have validated the enhanced capacity of SNPs to diagnose hybridity, particularly when compared with microsatellite loci (CamachoLSanchez et al., 2020; Sunde et al., 2020; Zimmerman et al., 2020; Szatmári et al., 2021). This is largely because microsatellite alleles are highly variable, with substantial homoplasy, and are thus rarely diagnostic (Estoup et al., 2000, and references therein). As such, they are less efficient either for determining hybridization or for assessing purity of core populations.

GADMA offered a finer-scale temporal perspective underlying our signal of historic introgression. The optimal demographic model supported periods of intermittent secondary contact/isolation between *S. catenatus* and *S. t. tergeminus*, likely mediated by glacial cycles and subsequent vicariant barriers. Results from PALEOTREE (Fig. 3A; Table 1) are consistent with the GADMA model, demonstrating that *S. catenatus* and *S. t. tergeminus* expanded northeast and southwest respectively, seemingly from an origin near the Great Plains contact zone.

The observed patterns of expansion are consistent with a recurring vicariant barrier, with potential candidates including the Mississippi and Missouri rivers (Szymanski, 1998; Sovic et al., 2016), which greatly increased in volume during glacial discharge from the Appalachian and Rocky mountains (respectively) during late Miocene [(∼4 Mya) (Bentley Sr et al., 2016)]. Periodic reductions in discharge then allowed dispersal northeastward via a ‘prairie corridor’ (Cook, 1992). Indeed, the Mississippi River has impacted phylogeographic patterns in other viperid snakes, such as *Agkistrodon* (Douglas et al., 2009) as well as *Sistrurus* (Sovic et al., 2016). In both cases, demographic modeling supported secondary contact consistent with our GADMA analysis, and with time scales similarly estimated (11 versu*s* 7 Kya) (Figs. 4A, 4B).

At a deeper temporal scale, the late Miocene/early Pliocene transition is also consistent with an estimated divergence of *S. catenatus* and *S. t. tergeminus* (∼4.86 Mya; 95% CI=4.09 – 6.25 Mya). The earliest known fossils attributable to *S. catenatus* or *S. tergeminus* date to the Pliocene in Kansas, Texas, and Nebraska, whereas the earliest identifiable *Sistrurus* date to the Miocene (∼9 Mya) in Nebraska. Thus, it is likely that the MRCA of *S. catenatus/S. tergeminus* originated in the Great Plains then dispersed both northeast and southwest, again consistent with our PALEOTREE results and the presence of a vicariant barrier. Northeastern *S. catenatus* could alternatively reflect newer, post-glacial populations that expanded from refugia, but in any case, our phylogeographic data indicate that the more ‘ancestral’ *S. t. tergeminus* lineages are Missouri and Iowa.

However, it is also possible that the observed introgression in Missouri individuals produced artificially-abbreviated nodal-root distances (Bangs et al., 2018). A similar pattern is found in *S. t. edwardsii,* where individuals with short nodal distances in Texas/New Mexico may represent artifacts of recent admixture in the southwestern contact zone. Although admixture seemingly affects diversification patterns in PALEOTREE, eastern Kansas does not contain admixed individuals and may thus represent the origin of both northeastern and southwestern diversifications, a prospect also consistent with the fossil record (Holman, 2000; Parmley and Holman, 2007).

The adaptive implications of the relictual hybrid lineage in Missouri are yet to be evaluated. Morphologically, intergraded features have been identified (Evans and Gloyd, 1948), but other attempts at classification using morphology, venom composition, and habitat utilization disagreed (Szymanski, 1998; Szymanski et al., 2016). Here, we note that hybrids between other rattlesnake taxa exhibit intermediate venom characteristics (Neri-Castro et al., 2022), but in this regard *Sistrurus* demonstrate high variability (Gibbs et al., 2009, 2011). Additional studies (e.g., whole-genome or transcriptome) that can potentially differentiate adaptations in the hybrid population would help to clarify taxonomic status, an important prospect if Missouri hybrids were to constitute a locally adapted hybrid lineage of potential conservation concern.

### 4.2. Recent divergence in the southwestern contact zone

Despite mixed ancestry in the southwestern contact zone (ADMIXTURE), introgression between *S. t. tergeminus* and *S. t. edwardsii* was not statistically supported by *D*-statistics, PHYLONETWORKS, and TREEMIX results (Fig. 6). Our results instead point to an ongoing divergence, as coupled with the observed paraphyletic relationship of the two *S. tergeminus* subspecies and their more recent divergence time. Our GADMA analyses concur (Figs. 4C, 4D), with strong, persistent, post-divergence migration and a lack of apparent vicariant barriers in the southwestern contact zone (Kubatko et al., 2011).

Although gene flow is ongoing between *S. t. tergeminus* and *S. t. edwardsii*, distinct ecological and physiological traits separate the two. For example, *S. t. edwardsii* prefers ectothermic prey (Holycross and Mackessy, 2002), and is found in xeric grasslands and dunes (Hammerson, 2000; Stebbins, 2003; Degenhardt et al., 2005; Wastell and Mackessy, 2011, 2016). On the other hand, *S. t. tergeminus* prefers mammalian prey (Holycross and Mackessy, 2002) and mesic prairies and grasslands (Seigel, 1986; Briggler and Johnson, 2021). These differences are reinforced by ecological niche modeling (Wooten and Gibbs, 2012), with temperature and precipitation regimes delineating the two. They also differ in venom composition, which is seemingly a corollary to their dietary preferences (but see Sanz et al., 2006; Gibbs and Mackessy, 2009; Gibbs et al., 2013).

The recent phylogeographic divergence estimated for *S. t. tergeminus* and *S. t. edwardsii* (1.44 Mya; 95% CI=1.26-1.73 Mya) coincides with the Pliocene-Pleistocene transition (∼1.5-2.0 Mya). The southwestern United States has since undergone many climatic and vegetational fluctuations (Savage, 1960; Findley, 1969; Morafka, 1977; Axelrod, 1983), oscillating between aridification/increasing xeric shrubs (Axelrod, 1979; Wilson and Pitts, 2010) *versus* the cooler climates and mesic habitat more suitable for *S. t. tergeminus* (MacKay and Elias, 1992; Pendall et al., 1999; Holycross, 2002; Holmgren et al., 2003; Wilson and Pitts, 2010). The region also seemingly arrived at its current state ∼8-4 Kya (Van Devender, 1977; MacKay and Elias, 1992; Hunter et al., 2001; Holmgren et al., 2003), again consistent with an expansion from ∼7 Kya, as modeled by GADMA (Fig. 4).

Importantly, given the scarcity of vicariant barriers in the region, recurring transitions between mesic and xeric habitat may have promoted long-term zones of mosaic contact during Pleistocene climate fluctuations, thus contributing to a gradual divergence between the two *S. tergeminus* ecotypes. A contemporary example of potential Pleistocene conditions can be found in northwestern Texas, where intermediate habitat provides a contact zone for numerous taxa (Swenson and Howard, 2005). Given the weak fossil record for *S. tergeminus* (Holman, 2000; Parmley and Holman, 2007), we cannot confirm if it was present in the southwest at the Pliocene-Pleistocene transition, or if it gradually encroached post-transition. Nevertheless, our observed diversification patterns, coupled with a recently formed but highly labile habitat, highlight the strong ecological component to its divergence, as noted in other taxa as well (Sartor et al., 2021).

### 4.3. *Conservation implications for* Sistrurus catenatus *and* S. t. tergeminus

Our analyses clearly delineate *S. catenatus* and *S. tergeminus,* and support previous assessments regarding their genetic diversity. First, GADMA showed a population bottleneck in *S. catenatus* consistent with recent estimates of a reduced *Ne* (Ochoa et al., 2020; Sovic et al., 2019), with habitat fragmentation as a major factor contributing to their decline (Szymanski et al., 2016). Second, the greater genetic diversity we noted in *S. t. tergeminus* is also consistent with other recent studies that identified a greater *Ne* when compared to *S. catenatus* (Sovic et al., 2016). The *S. t. tergeminus* habitat is less affected by anthropogenic fragmentation in (at least) parts of its range (Greene, 1997; Szymanski, 1998; McCluskey and Bender, 2015), although our previously undescribed population structure warrants further investigation.

The ADMIXTURE results for *S. t. tergeminus* recognized two populations as distinct: Northeastern (Missouri/eastern Kansas) and southwestern (southcentral Kansas/Oklahoma/Texas) (Fig. 3B). Although reproductive boundaries between the two are seemingly porous, their primary separation corresponds to the Arkansas River, a barrier likewise found as significant in other studies (Fontanella et al., 2008; Ruane et al., 2014; Herman and Bouzat, 2016). Differentiation of central Texas and Oklahoma has also been previously noted (Ryberg et al., 2015), with our study again in agreement. However, our results also contrast with earlier studies that found minimal structure within *S. t. tergeminus* (McCluskey and Bender, 2015). The latter study was geographically constrained to KS and MO, and thus may have missed the southern population. Earlier studies also failed to support subspecific boundaries (Ryberg et al., 2015; Bylsma et al., 2021). The results from Ryberg et al. (2015) were based on microsatellite loci or mitochondrial DNA and intron sequences, and thus may have had a reduced capacity to discriminate relative to our large SNP dataset (Rašić et al., 2014; Vendrami et al., 2017), particularly given the complications that can be introduced by admixture (Haasl and Payseur, 2011). Finally, we disagree with Bylsma et al. (2021) with regard to *S. tergeminus* as a monotypic entity, and note that subspecific divergence, while not discrete, is also apparent in their PCA results. It may be that the greater number of markers in our study [N=10,190 versus N=171 (Bylsma et al., 2021)] was better able to detect fine-scale population structure.

### 4.4 Conservation implications for Sistrurus tergeminus edwardsii

The GADMA results are concordant with previous research indicating that *S. t. edwardsii* has undergone recent population growth (Anderson et al., 2009). However, although *S. t. edwardsii* seemingly has elevated genetic diversity, its habitat is either disappearing or undergoing seriously fragmentation (Lowe et al., 1986; Greene, 1997; Werler and Dixon, 2010), which in turn implies an impending ‘drift debt’ situation (per *S. catenatus*; Ochoa et al., 2020) and reflects the time lag between habitat fragmentation and a decline in genetic diversity. The fine-scale population structure detected in *S. t. edwardsii* from Texas/New Mexico/Arizona and Colorado further support the potential of a drift-debt hypothesis, in concordance with their topographic disjunction and as an addendum to previous research (e.g., Mackessy, 2005; Anderson et al., 2009). It also underscores the need for further conservation genetic studies at a finer scale, particularly given the probability that the southwestern *Sistrurus* contact zone may shift northward with oncoming climate change (Walkup et al., 2022). In this sense, recognizing the extent of existing genetic diversity may help not only to predict their future evolutionary trajectory but also conserve their current one.

### 4.5 Taxonomic implications

*Sistrurus t*. *tergeminus* and *S. t. edwardsii* are clearly less divergent than are *S. catenatus* and *S. tergeminus* ssp. Nonetheless, while the genetic barriers between both pairs are relatively porous at their contact zones (northeast and southwest), distinct population structure is apparent within and among subspecies. This should establish each as worthy of conservation consideration. Furthermore, the hybridization and introgression observed in the two contact zones is of potential evolutionary significance, but at disparate temporal scales. The disjunct Missouri populations are potentially locally adapted and relictual, stemming from an ancient introgression event. The issue of ancestral hybridization has been previously recognized as yielding evolutionarily distinct lineages, while at the same time provoking taxonomic as well as management issues (Pacheco-Sierra et al., 2018). Similarly, *S. t. tergeminus* and *S. t. edwardsii* in the southwest may also be undergoing a primary (and more contemporaneous) divergence event.

Taxonomic considerations provide a very useful model for biodiversity, but need not be applied in every managerial situation because they do not always juxtapose with the needs of species with complex demographic histories. For example, hybridization is a long-standing biological phenomenon that has repeatedly confounded classificatory applications (Fitzpatrick et al., 2015). Yet in ambiguous situations (as herein), it often becomes the recipient of a ‘taxonomically-based’ managerial decision. Despite the now-recognized prevalence of hybridization in nature (Taylor and Larson, 2019), the ‘one-shoe-fits-all’ application of taxonomy becomes a convenient hook from which to hang discordant events (Dussex et al., 2018). The negative connotations surrounding ‘hybridity’ – i.e., as a major contributor to extinction – is an inherent, historic, and a sufficient superficial reason to do so (Draper et al., 2021).

### 4.6 Application under ‘New Conservation’

Rather, a more nuanced path is clearly needed for administrators, managers, and applied biologists (Jackiw et al., 2015), one that has been repeatedly driven home by contemporary results using highly-sophisticated molecular techniques and nuanced but powerful analytic approaches (Bohling, 2016). Simply put, legislative goals and scientific classification are not persistent handmaidens for one another, and conservation biology must understand and react accordingly.

Some of this ambiguity stems from the scientific literature on hybridization, particularly regarding the ‘how’s and why’s’ of hybrid protection. Earlier viewpoints interpreted the biological basis of hybridity as a barometer, i.e., occurring within a natural versus anthropogenic context (Allendorf et al., 2001). This approach, while germane for its time, is now inadequate as viewed from a more contemporary and anthropogenic perspective, which seeks to avoid value-loaded terms such as ‘naturalness’ as a guiding principle for hybrid management (Donfrancesco and Luque-Lora, 2021). In fact, anthropogenic hybridization is now viewed as having positive conservation considerations (Chan et al., 2019; Ottenburghs, 2021), and has already been utilized as a managerial strategy (Zecherle et al., 2021).

The tenor of this debate parallels an earlier one, also separated by 20 years, focusing on how conservation should now be defined. ‘New’ conservation (Kareiva and Marvier, 2012) suggests that humans are the dominant ecological force on the planet and can no longer remain separated from nature, with conservation achievements gauged in large part by its anthropogenic relevance. Legacy perspectives (Soulé, 2013) were deemed as strictly biological (rather than human-oriented), with narrower diagnoses and unrealistic solutions incompatible with an irreversible Anthropocene. Here, the preservation and/or restoration of past ecosystems is deemed a losing proposition. However, the debate eventually dwindled. Each viewpoint is complementary in its desire to sustain nature, albeit with different philosophies and approaches. Hopefully parallel arguments regarding hybrid conservation and management instead reach an operational consensus rather that a ‘fade-to-gray.’

Clearly the admixed species-pairs (discussed above) warrant federal protection. We base this upon their genetic differentiation, as well as their aforementioned ecological and physiological disparities (section 4.2). We support our contention based on four aspects that should underlie the protection of hybrid populations (vonHoldt et al., 2018): (1) One (or both) of the parentals is endangered; (2) Hybridization is not an intentional cross-breeding situation, unless it is a conservation efforts to introduce needed genetic heterogeneity; (3) Frequency of hybridization is either low, or has occurred far enough in the past such that the majority of individuals are hybrid genotypes; and (4) Failure to protect would likely result in the loss of most or all of the alleles that distinguish the endangered parental type(s).

At a minimum, both components of the species-pair should be recognized as ESUs. The population structure within each may also reflect a cryptic genetic diversity potentially warranting consideration as MUs.

### 4.6 Conclusions

Herein, we have expanded upon and contextualized the population structure and phylogeography of *S. catenatus* and *S. tergeminus* ssp. In doing so, we characterize contrasting biogeographic processes underlying two North American contact zones and demonstrated that constituent taxa have a more complex and diversified evolutionary history than previously surmised. In the northeastern *S. tergeminus* range, speciation has been strongly influenced by vicariant barriers and secondary contact, whereas in the southwest, a primary divergence event is seemingly now underway, as facilitated by ecological and physiological differences and a habitat whose availability has fluctuated since the Pliocene. The two contact zones highlight the substantial direct and indirect impacts that emerge when regional biogeography intersects with climate change across closely related biodiversity elements. In this sense, numerous (and relatively cryptic) biodiversity transitions that have occurred prior to the Pleistocene/Holocene boundary (per ancient DNA investigations; Cooper et al., 2015) reinforce the importance of climate change in driving population structure, and ultimately impacting population declines and extinctions. It also provides an excellent ongoing reference for the study of biogeography and genetic diversity across similar biodiversity elements arrayed within and among nearby regions.

## Authorship contribution statement

**Bradley T. Martin:** Conceptualization, Methodology, Software, Formal analysis, Investigation, Writing – Original Draft, Visualization. **Marlis R. Douglas:** Conceptualization, Resources, Data Curation, Writing – Review & Editing, Supervision, Project administration, Funding acquisition. **Tyler K. Chafin:** Methodology, Software, Formal Analysis, Writing – Review & Editing, Visualization. **John S. Placyk, Jr.:** Conceptualization, Resources, Writing – Review & Editing. **Steven P. Mackessy:** Conceptualization, Resources, Writing – Review & Editing. **Jeffrey T. Briggler:** Conceptualization, Resources, Writing – Review & Editing. **Michael E. Douglas:** Conceptualization, Resources, Data Curation, Writing – Review & Editing, Supervision, Project administration, Funding acquisition.

## Declaration of Competing Interest

The authors declare that they have no known competing financial interests or personal relationships.

## Data availability statement

Raw ddRADseq data are available as a Sequence Read Archive (SRA) on the GenBank Nucleotide Database [Accession Nos. provided upon acceptance]. All relevant input, output, and configuration files are available in an Open Science Framework (OSF) data repository [DOI provided upon acceptance]. Any custom scripts used in the analyses are available in GitHub repositories [https://github.com/btmartin721/ddrad_scripts, https://github.com/tkchafin/scripts, https://github.com/tkchafin/makeCompD].

## Acknowledgments

This manuscript represents one component of materials submitted by BTM in partial fulfillment of the Ph.D. degree in Biological Sciences at University of Arkansas, Fayetteville. We thank the countless colleagues, volunteers, museums, and agencies who have either collected or contributed samples for this project (Supplementary Table S1). We also thank several members of the Douglas Lab and University of Arkansas faculty for their invaluable guidance, to include: A. Alverson, W. Anthonysamy, M. Bangs, J. Beaulieu, S. Mussmann, J. Pummill, and Z. Zbinden. This work was supported by University of Arkansas endowments to MRD and MED, and an NSF Postdoctoral Research Fellowship in Biology to TKC (DBI: 2010774). The Arkansas High Performance Computing Cluster (AHPCC) and Jetstream cloud (XSEDE #TG-BIO160065) provided computational resources. The findings, conclusions, and opinions expressed in this article represent those of the authors and do not necessarily represent the views of the NSF nor other affiliated or contributing organizations.

Appendix A. Supplementary material

Supplementary data to this article can be found online at [URL to be supplied by journal].

## SUPPLEMENTAL TABLES AND FIGURES

**Table S1:** Sample/ collector data for *N*=226 *Sistrurus*, *Crotalus*, and *Agkistrodon* individuals.

| ID | Date | Sex | Collector(s)/ Source(s) | State | County |
| --- | --- | --- | --- | --- | --- |
| SMAR1 | X-00 | F | C. Montgomery | AR | Washington |
| SMAR4 | X-00 | M | K. Irwin | AR | Montgomery |
| SMAR5 | 8-VII-00 | F | K. Irwin | AR | Marion |
| CM2 |  | M | A. Holycross | AZ | Coconino |
| CMO72 | 04-VII-02 | M | G. Carpenter | AZ | Mohave |
| CS8 | 07-VIII-00 |  | G. Schuett | AZ | Yavapai |
| CVC32 | VIII-98 | - | C. Meachum | AZ | Pima |
| CVC6 |  | M | G. Schuett | AZ | Coconino |
| CWW6 | VII-93 |  | T. LaDuc | AZ | Cochise |
| CWW7 | VIII-02 |  | G.W. Schuett | AZ | Cochise |
| SCE10 |  | F | Phx. Zoo | AZ | Cochise |
| SCE11 |  | F | Phx. Zoo | AZ | Cochise |
| SCE12 |  | F | Phx. Zoo | AZ | Cochise |
| SCE9 |  | M | Phx. Zoo | AZ | Cochise |
| SICA52G |  |  | L. Gibbs; A. Holycross | AZ | Cochise |
| SICA53G |  |  | L. Gibbs; A. Holycross | AZ | Cochise |
| SICA54G |  |  | L. Gibbs; A. Holycross | AZ | Cochise |
| SCCO1 |  | F | S. Mackessy | CO | Lincoln |
| SCCO10 |  |  | S. Mackessy | CO | Lincoln |
| SCCO12 |  |  | S. Mackessy | CO | Lincoln |
| SCCO13 |  |  | S. Mackessy | CO | Lincoln |
| SCCO14 |  |  | S. Mackessy | CO | Lincoln |
| SCCO15 |  |  | S. Mackessy | CO | Lincoln |
| SCCO16 |  |  | S. Mackessy | CO | Lincoln |
| SCCO17 |  |  | S. Mackessy | CO | Lincoln |
| SCCO18 |  |  | S. Mackessy | CO | Lincoln |
| SCCO19 |  |  | S. Mackessy | CO | Lincoln |
| SCCO2 |  | F | S. Mackessy | CO | Lincoln |
| SCCO20 |  |  | S. Mackessy | CO | Lincoln |
| SCCO21 |  |  | S. Mackessy | CO | Lincoln |
| SCCO22 |  |  | S. Mackessy | CO | Lincoln |
| SCCO23 |  |  | S. Mackessy | CO | Lincoln |
| SCCO24 |  |  | S. Mackessy | CO | Lincoln |
| SCCO25 |  |  | S. Mackessy | CO | Lincoln |
| SCCO26 |  |  | S. Mackessy | CO | Lincoln |
| SCCO27 |  |  | S. Mackessy | CO | Lincoln |

**Table S1 (Cont.)**
| **ID** | **Date** | **Sex** | **Collector(s)/ Source(s)** | **State** | **County** |
| --- | --- | --- | --- | --- | --- |
| SCCO28 |  |  | S. Mackessy | CO | Lincoln |
| SCCO29 |  |  | S. Mackessy | CO | Lincoln |
| SCCO3 |  | M | S. Mackessy | CO | Lincoln |
| SCCO35 |  |  | S. Mackessy | CO | Lincoln |
| SCCO37 |  |  | S. Mackessy | CO | Lincoln |
| SCCO38 |  |  | S. Mackessy | CO | Lincoln |
| SCCO39 |  |  | S. Mackessy | CO | Lincoln |
| SCCO4 |  |  | S. Mackessy | CO | Lincoln |
| SCCO41 |  |  | S. Mackessy | CO | Lincoln |
| SCCO42 |  |  | S. Mackessy | CO | Lincoln |
| SCCO43 |  |  | S. Mackessy | CO | Lincoln |
| SCCO44 |  |  | S. Mackessy | CO | Lincoln |
| SCCO45 |  |  | S. Mackessy | CO | Lincoln |
| SCCO47 |  |  | S. Mackessy | CO |  |
| SCCO48 |  |  | S. Mackessy | CO | Baca |
| SCCO49 |  |  | S. Mackessy | CO | Baca |
| SCCO5 |  |  | S. Mackessy | CO | Lincoln |
| SCCO50 |  |  | S. Mackessy | CO | Cheyenne |
| SCCO6 |  |  | S. Mackessy | CO | Lincoln |
| SCCO9 |  |  | S. Mackessy | CO | Lincoln |
| APFL1 |  |  | W. Hayes | FL | Liberty |
| SMFL12 | 21-VIII-03 |  | P. Moler | FL | Wakulla |
| SMFL13 | 21-VIII-03 |  | P. Moler | FL | Okaloosa |
| SMFL5 |  |  | S. Conners | FL | Collier |
| SMFL6 | 23-XI-01 |  | S. Conners | FL | Miami-Dade |
| SMFL8 | 27-XII-02 |  | G. Pyron | FL | Franklin |
| SMGA1 | 16-IV-03 |  | G. Schuett | GA | Cherokee |
| SMGA2 | 15-VI-03 | M | C. Ponder | GA | Ware |
| SCIA3 | 20-IV-04 | F | T. VanDeWalle | IA | Bremer |
| SCIA4 | 3-V-04 | M | T. VanDeWalle | IA | Bremer |
| SCIA5 | 3-V-04 | F | T. VanDeWalle | IA | Bremer |
| SCIA7 | 3-V-04 | F | T. VanDeWalle | IA | Bremer |
| SCIA8 | VIII-90 |  | J. Christiansen | IA | Scott |
| SCIL1 |  | - | C. Phillips | IL | Clinton |
| SCIL2 |  | - | C. Phillips | IL | Clinton |
| SCIL3 |  | - | C. Phillips | IL | Clinton |
| SCIL4 | 08-VI-01 | M | T. Anton | IL | Cook |
| SCIL6 |  | F |  | IL | Carlisle |
| SCIL7 |  | F |  | IL | Carlisle |
| SCIL8 |  | F |  | IL | Carlisle |
| ACKS3 | VI-00 |  | H. Alamillo | KS | Johnson |

**Table S1 (Cont.)**
| **ID** | **Date** | **Sex** | **Collector(s)/ Source(s)** | **State** | **County** |
| --- | --- | --- | --- | --- | --- |
| ACKS5 | 10-V-00 |  | C. Shiel | KS | Douglas |
| SCKS1 |  | F | S. Mackessy | KS | Barton |
| SCKS11 | 19-IX-04 |  | T. Taggart | KS | Clark |
| SCKS12 | 20-IX-04 |  | T. Taggart | KS | Comanche |
| SCKS13 | 20-IX-04 |  | C.J. Schmidt | KS | Kiowa |
| SCKS14 |  |  |  | KS | Chase |
| SCKS15 |  |  |  | KS | Chase |
| SCKS16 |  |  |  | KS | Barber |
| SCKS17 |  |  |  | KS | Chase |
| SCKS18 |  |  |  | KS | Kiowa |
| SCKS19 |  |  |  | KS | Comanche |
| SCKS2 |  | M | S. Mackessy | KS | Barton |
| SCKS20 |  |  |  | KS | Allen |
| SCKS21 |  |  |  | KS | Barber |
| SCKS22 |  |  |  | KS | Reno |
| SCKS23 |  |  |  | KS | Russell |
| SCKS24 |  |  |  | KS | Kiowa |
| SCKS25 |  |  |  | KS | Douglas |
| SCKS26 |  |  |  | KS | Stafford |
| SCKS3 |  | M | S. Mackessy | KS | Barton |
| SCKS4 |  | M | S. Mackessy | KS | Barton |
| SCKS5 |  | M | S. Mackessy | KS | Barton |
| SCKS6 | 28-IV-01 |  | D. Fogell | KS | Pottawatomie |
| SCKS7 | VI-02 | F | D. Fogell | KS | Chase |
| SCKS8 |  |  | D. Shepard | KS | Butler |
| SCKS9 | 23-VI-04 |  | J. Voelkler | KS | Washington |
| SICA18G | 25-IV-09 |  | R. Brown, KU, P. Ingram | KS | Chase |
| SICA21G |  |  | B. coyner SNOMNH | KS | Butler |
| SICA27G | 15-X-04 |  | C.J. Schmidt SMNH, M. Washburne | KS | Chautauque |
| E058 |  |  | J. Moore | MI |  |
| E114 |  |  | J. Moore | MI |  |
| E270 |  |  | J. Moore | MI |  |
| E294 |  |  | J. Moore | MI |  |
| E301 |  |  | J. Moore | MI |  |
| E356 |  |  | J. Moore | MI |  |
| E513 |  |  | J. Moore | MI |  |
| E545 |  |  | J. Moore | MI |  |
| E577 |  |  | J. Moore | MI |  |
| E635 |  |  | J. Moore | MI |  |
| E794 |  |  | J. Moore | MI |  |
| E819 |  |  | J. Moore | MI |  |

**Table S1 (Cont.)**
| **ID** | **Date** | **Sex** | **Collector(s)/ Source(s)** | **State** | **County** |
| --- | --- | --- | --- | --- | --- |
| SCMI2 | VI-01 |  | K. Schuett | MI | Hillsdale |
| SICA46G |  |  | J. Moore | MI |  |
| SCMO1 | 26-IX-01 | F | F. Durbian | MO | Holt |
| SCMO11 | VI-02 | F |  | MO | Chariton |
| SCMO12 | VI-02 | M |  | MO | Chariton |
| SCMO13 | VI-02 |  |  | MO | Chariton |
| SCMO14 | 02-IV-05 | F | R.Seigel/T.Crabill | MO | Linn |
| SCMO15 | 02-IV-05 | M | R.Seigel/T.Crabill | MO | Linn |
| SCMO16 | 02-IV-05 | F | R.Seigel/T.Crabill | MO | Linn |
| SCMO17 | 03-IV-05 | M | R.Seigel/T.Crabill | MO | Linn |
| SCMO18 | 03-IV-05 | M | R.Seigel/T.Crabill | MO | Linn |
| SCMO19 | 03-IV-05 | F | R.Seigel/T.Crabill | MO | Linn |
| SCMO2 | 27-IX-01 | M | F. Durbian | MO | Holt |
| SCMO20 | 04-IV-05 | F | R.Seigel/T.Crabill | MO | Linn |
| SCMO21 | 04-IV-05 | M | R.Seigel/T.Crabill | MO | Linn |
| SCMO22 | 04-IV-05 |  | R.Seigel/T.Crabill | MO | Linn |
| SCMO23 | 04-IV-05 |  | R.Seigel/T.Crabill | MO | Linn |
| SCMO24 |  |  |  | MO | Chariton |
| SCMO25 |  |  |  | MO | Linn |
| SCMO3 | 27-IX-01 | M | F. Durbian | MO | Holt |
| SCMO4 | 27-IX-01 | M | F. Durbian | MO | Holt |
| SCMO5 | 27-IX-01 | F | F. Durbian | MO | Holt |
| SCMO6 | 27-IX-01 | M | F. Durbian | MO | Holt |
| SCMO7 | 02-X-01 | F | F. Durbian | MO | Holt |
| SCMO8 | 02-X-01 | F | F. Durbian | MO | Holt |
| SCMO9 | VI-02 | J |  | MO | Chariton |
| SMMS1 | VI-01 | F | VanDevender | MS | Copiah |
| SCNE1 | VIII-00 | F | D. Fogell | NE | Pawnee |
| SCNE2 | VIII-00 | M | D. Fogell | NE | Pawnee |
| SCNE3 | VIII-00 | F | D. Fogell | NE | Pawnee |
| SCNE4 | V-99 |  | M. Ingrasci | NE | Russell |
| CVV12 | 09-IV-00 | F | C. Painter | NM | Chaves |
| SCE1 | 22-VII-00 | J | D.&K.Salceies | NM | Bernalillo |
| SCE2 | 14-VII-00 | J | D.&K.Salceies | NM | Bernalillo |
| SCE3 | 7-VII-00 | J | D.&K.Salceies | NM | Bernalillo |
| SCE4 | 3-VII-00 | J | D.&K.Salceies | NM | Bernalillo |
| SCE5 | 22-VFII-00 | J | D.&K.Salceies | NM | Bernalillo |
| SCE6 | 4-VII-00 | J | D.&K.Salceies | NM | Bernalillo |
| SCE7 | 14-VII-00 | J | D.&K.Salceies | NM | Bernalillo |
| SCE8 | IX-00 | M | C. Painter | NM | Socorro |
| SCNM1 | 29-VI-00 | M | T. LaDuc | NM | Valencia |

**Table S1 (Cont.)**
| **ID** | **Date** | **Sex** | **Collector(s)/ Source(s)** | **State** | **County** |
| --- | --- | --- | --- | --- | --- |
| SCNM2 | 06-X-96 |  | T.R. Jones | NM | Lincoln |
| SCNM3 | 18-V-98 |  | L. Kamees | NM | Socorro |
| SCNM4 |  |  | B. Christman | NM | Otero |
| SCNM5 | VII-01 |  | B. Christman | NM | Otero |
| SCNM6 |  |  | B. Mackin | NM | Grant |
| SCNM7 |  |  | B. Mackin | NM | Grant |
| SICA67G |  |  | Elda Sanchez NNTRC | NM | Otero |
| SCNY1 | 09-VI-92 |  | G. Johnson | NY | Monroe |
| SCNY2 | 09-VI-92 |  | G. Johnson | NY | Monroe |
| SCNY3 | 4-V-92 |  | G. Johnson | NY | Onandaga |
| SCNY4 | 18-VI-92 | F | G. Johnson | NY | Onandaga |
| SCOH1 | 25-VIII-00 | - | M. Spille | OH | Greene |
| SCOH11 |  |  | M. Spille | OH | Greene |
| SCOH12 |  |  | M. Spille | OH | Greene |
| SCOH16 | 29-V-92 | F | G. Johnson | OH | Clark |
| SCOH17 | 29-V-92 | F | G. Johnson | OH | Clark |
| SCOH3 | 25-VIII-00 | - | M. Spille | OH | Greene |
| SCOH8 | 06-IX-00 | - | D. Wynn | OH | Wyandot |
| SCOK08 |  |  | B. coyner SNOMNH | OK | Roger Mills |
| SCOK1 | 26VIII05 | F | D.Shepard/L.Vitt | OK | Beckham |
| SCOK10 |  |  |  | OK | Dewey |
| SCOK11 |  |  |  | OK | Beckham |
| SCOK12 |  |  |  | OK | Rogers |
| SCOK2 | 26VIII05 | J | D.Shepard/L.Vitt | OK | Beckham |
| SCOK3 |  |  | D.Shepard/L.Vitt | OK | Roger Mills |
| SCOK4 |  |  | D.Shepard/L.Vitt | OK | Roger Mills |
| SCOK5 |  |  | D.Shepard/L.Vitt | OK | Ellis |
| SCOK6 |  |  | D.Shepard/L.Vitt | OK | Blaine |
| SICA65G |  |  | Elda Sanchez NNTRC | OK | Comanche |
| SMOK3 | 7-VII-05 |  | C. Whitney | OK | Tulsa |
| SICA57G |  |  | L. Gibbs | ON | Dorcas Bay |
| SMSC1 | 15-XI-00 | F | B. Starrett | SC | Charleston |
| APTX1 |  |  | W. Hayes | TX | Eastland |
| CSTX6 | 2003 |  | B. Mackin | TX | Culberson |
| CSTX6 | 2003 |  | B. Mackin | TX | Culberson |
| SCT1 | 9-V-00 | - | K. McCoy | TX | Concho |
| SCT2 | 24-VI-00 | M | T. Hibbitts | TX | Parker |
| SCT3 | 24-VI-00 | M | T. Hibbitts | TX | Hood |
| SCTX1 |  | M | A. Price | TX | Cottle |
| SCTX10 |  |  |  | TX | Cottle |
| SCTX11 |  |  |  | TX | Cottle |

**Table S1 (Cont.)**
| **ID** | **Date** | **Sex** | **Collector(s)/ Source(s)** | **State** | **County** |
| --- | --- | --- | --- | --- | --- |
| SCTX14 |  |  |  | TX | Runnels |
| SCTX15 |  |  |  | TX | Motley |
| SCTX16 |  |  |  | TX | Dickens |
| SCTX17 |  |  |  | TX | Borden |
| SCTX18 |  |  |  | TX | Cottle |
| SCTX2 | 29-V-01 |  | T. LaDuc | TX | Andrews |
| SCTX3 | 06-V-01 |  | T. LaDuc | TX | Crockett |
| SCTX4 | 26-V-01 |  | T. LaDuc | TX | Motley |
| SCTX5 | 26-VI-01 |  | T. LaDuc | TX | Crockett |
| SCTX8 | 114-IX-99 | J | Tamu-K | TX | Galveston |
| SCTX9 |  |  | GladysPorterZoo | TX | Starr |
| SICA3G | 9-IX-14 |  | Matador WMA, S. Hein, M. Barazowski | TX | Cottle |
| SICA48G |  |  | Matador WMA, S. Hein, M. Barazowski | TX | Cottle |
| SICA61G | 8-VI-15 |  | R. Couvillian | TX | Jim Hogg |
| SICA62G | 26-VI-15 |  | S. Hein and S. Pitts | TX | Ward |
| SICA66G |  |  | Elda Sanchez NNTRC | TX | Nueces |
| SICA70G |  |  | T. Hibbitts | TX | Archer |
| SMTX1 | 31-VII-06 |  | T. Sinclair | TX | Montgomery |
| SMTX2 | 31-VII-06 |  | T. Sinclair | TX | Montgomery |
| SCWI10 | 10-X-01 | M | E. McCumber | WI | Buffalo |
| SCWI2 | 01-VIII-00 | F | via ATH | WI | Buffalo |
| SCWI6 | 30-VIII-01 | F | E. McCumber | WI | Buffalo |
| SCWI7 | 02-IX-01 | F | E. McCumber | WI | Buffalo |
| SCWI8 | 02-IX-01 | M | E. McCumber | WI | Buffalo |
| SCWI9 | 12-VII-01 | M | E. McCumber | WI | Buffalo |
| CVV56 |  |  | W. Hayes | WY | Sheridan |

**Table S2:** Number of sequenced individuals per *Sistrurus* taxon.

| **Taxonomic ID** | **Count** |
| --- | --- |
| *S. catenatus* | 44 |
| *S. tergeminus tergeminus* | 85 |
| *S. tergeminus edwardsii* | 68 |
| *S. miliarius miliarius* | 2 |
| *S. miliarius barbouri* | 6 |
| *S. miliarius streckeri* | 7 |
| **Total** | 212 |

**Figure S1:**
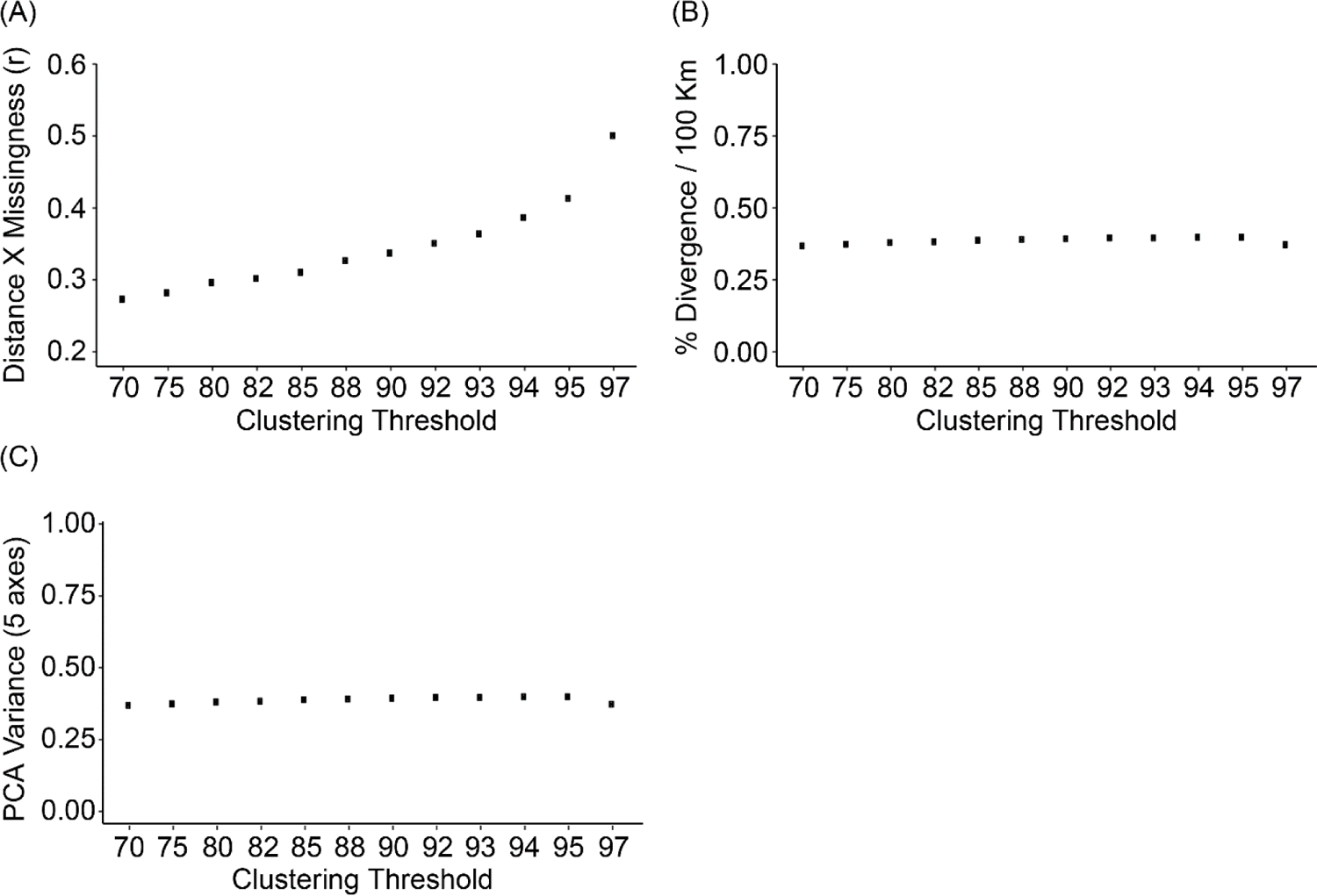
Assessments of varying IPYRAD clustering threshold parameters applied to *Sistrurus* ddRAD data. (A) Pearson’s correlation coefficient (r) between genetic distance and the percentage of missing data in the alignment. (B) Percent sequence divergence per 100 kilometers (Km) as a measure of IBD (isolation by distance); (C) principal component analysis (PCA) variance across five principal component axes.

**Figure S2:**
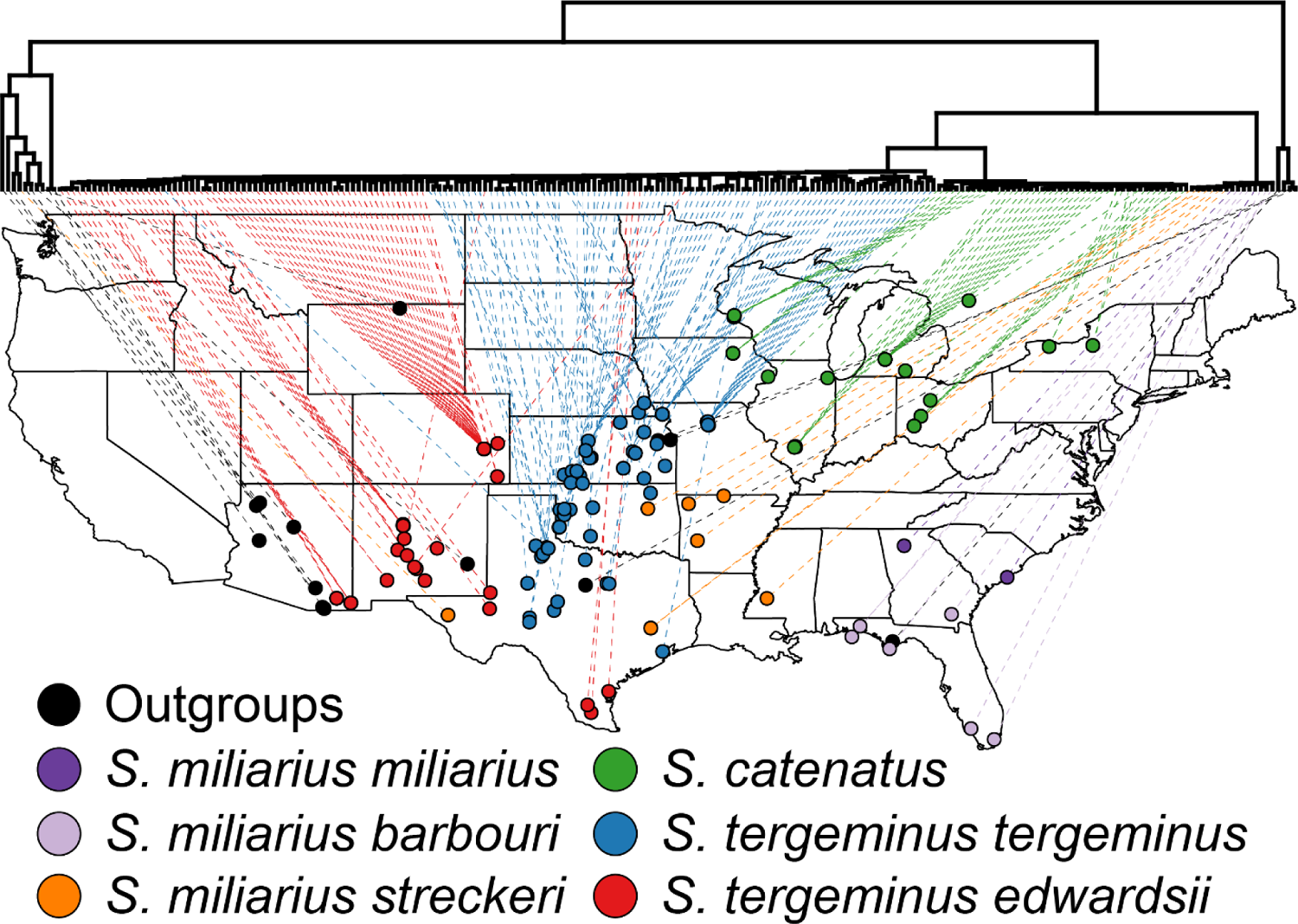
IQ-TREE phylogeny depicting the geographic localities per tip, color-coded to each *Sistrurus* species and subspecies (per field identification). Outgroups include several species of *Crotalus* and *Agkistrodon*. Dashed lines link tips with the corresponding mapped sample.

**Figure S3:**
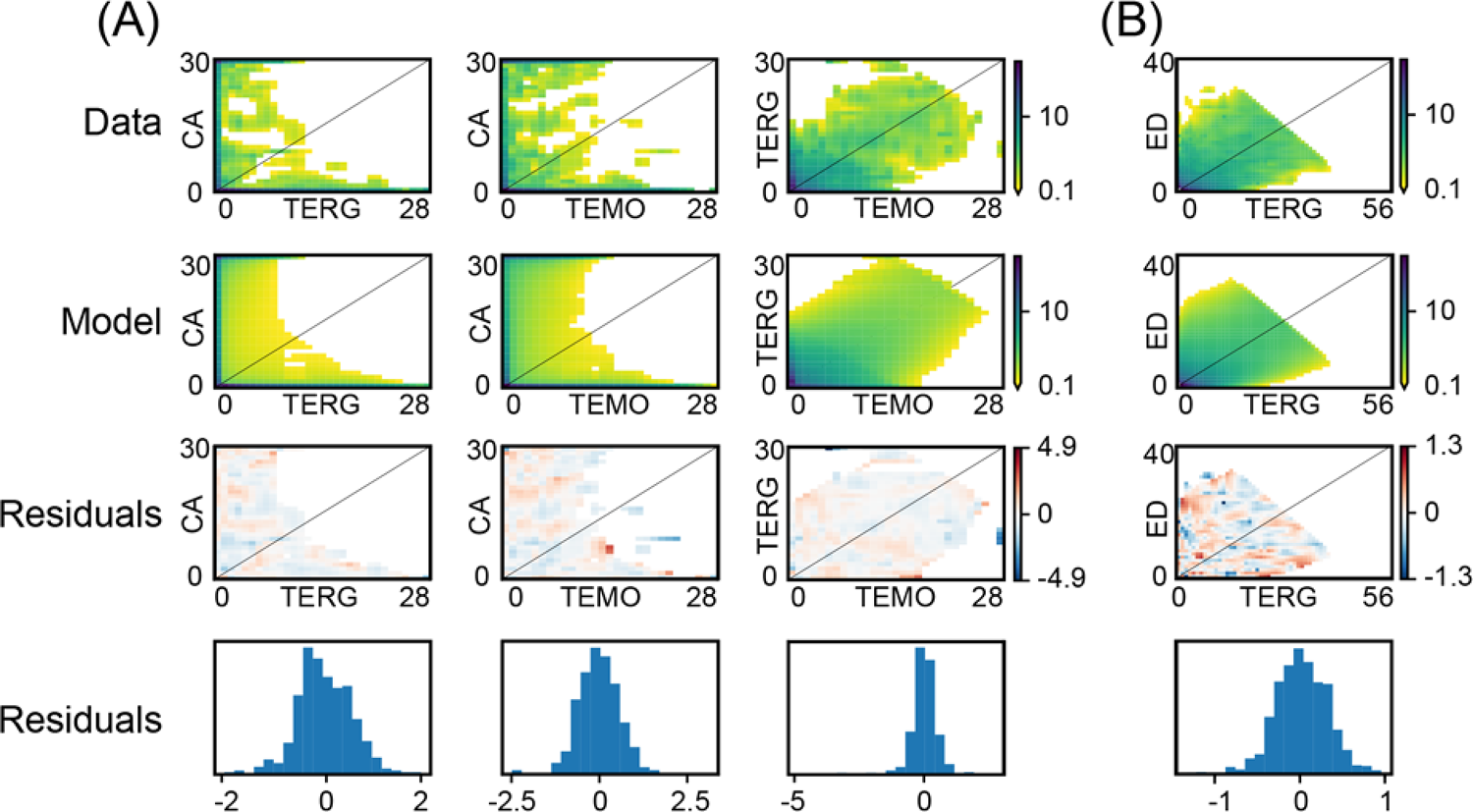
S*i*strurus allele frequency spectra (AFS) for the GADMA data, optimal model, and residuals as heatmaps and histograms. Pairwise AFS plots represent the following populations: (A) CA=*S. catenatus*, TEMO=*S. tergeminus tergeminus* from Missouri, and TERG=*S. t. tergeminus* from all other localities; (B) TERG=*S. t. tergeminus* from all localities except Missouri and ED=*S. tergeminus edwardsii*.

**Figure S4:**
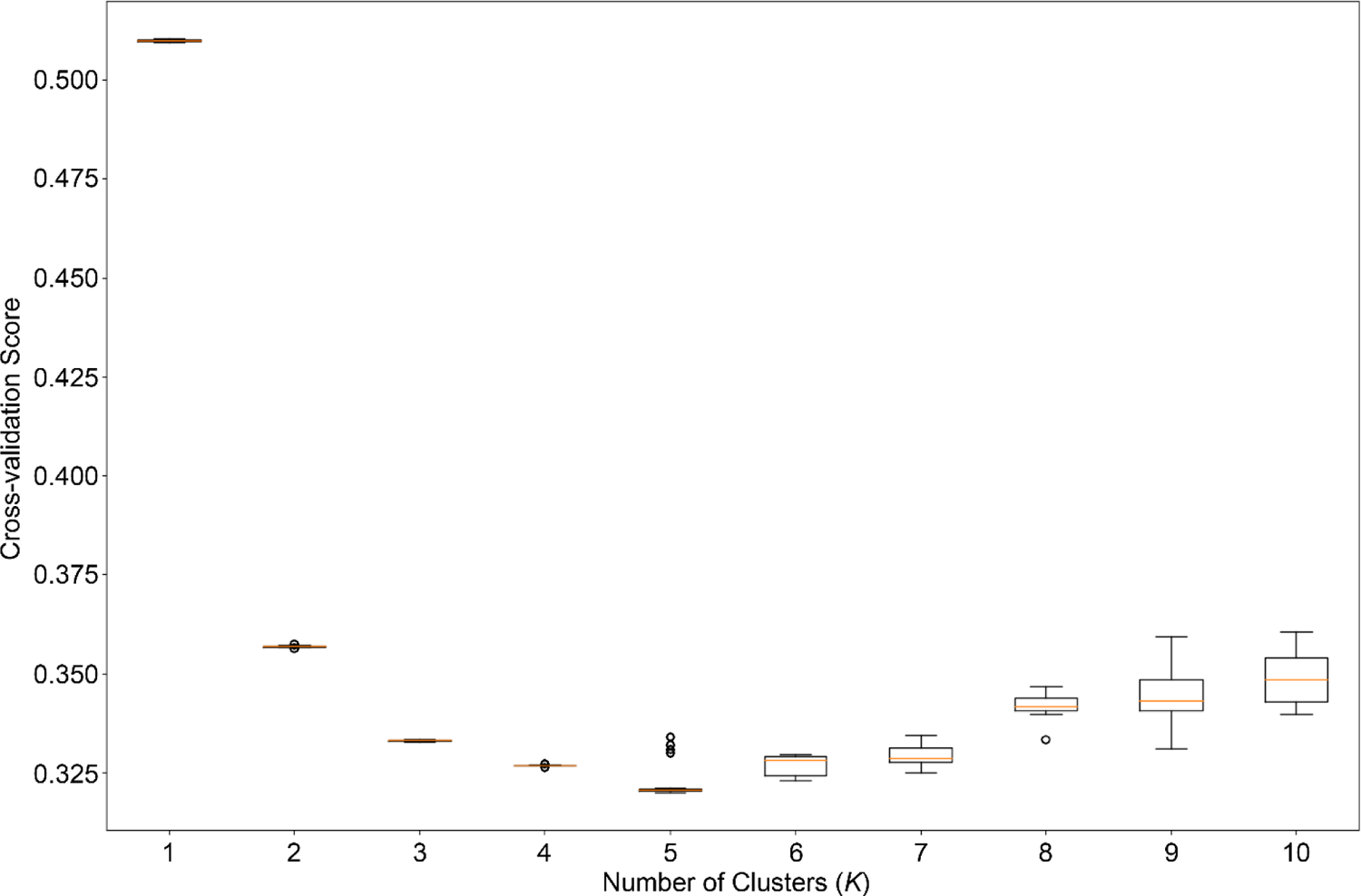
Boxplots for cross-validation scores from an ADMIXTURE analysis between *Sistrurus catenatus*, *S. tergeminus tergeminus*, and *S. t. edwardsii*. ADMIXTURE was run with 20 replicates per *K* and 20-fold cross-validation.

**Figure S5:**
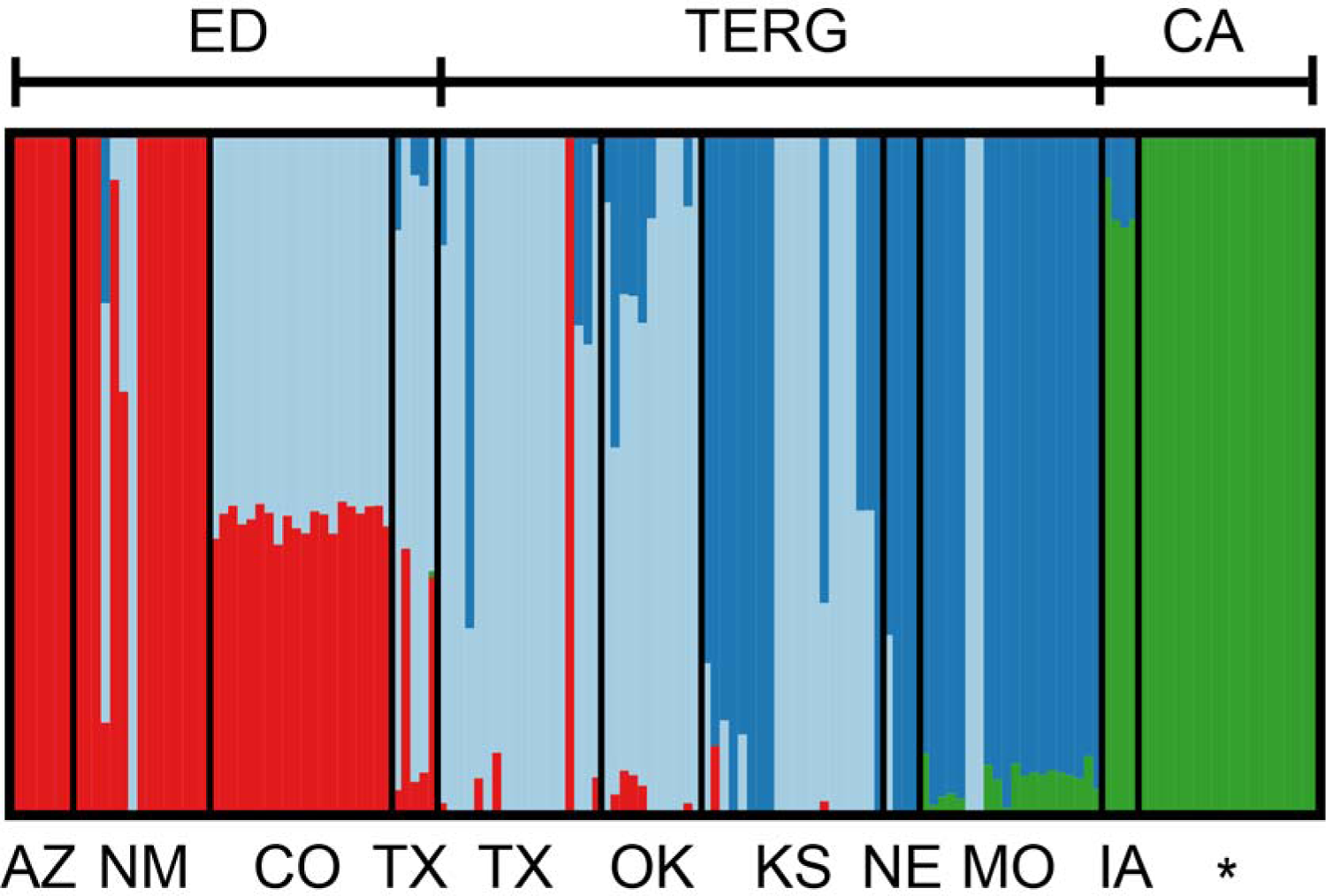
ADMIXTURE bar-plot depicting proportions among *K*=4 *Sistrurus* populations. Each bar represents one individual and those with mixed colors indicate mixed ancestry. The top row indicates subspecies designation as identified in the field: ED=*S. tergeminus edwardsii*, TERG=*S. tergeminus tergeminus*, CA=*S. catenatus*. The barplot is further partitioned into sub-populations based on U.S. state locality (IA=Iowa, MO=Missouri, NE=Nebraska, KS=Kansas, OK=Oklahoma, TX=Texas, CO=Colorado, NM=New Mexico, AZ=Arizona). The asterisk for *S. catenatus* indicates that individuals were included from multiple state localities.

**Figure S6:**
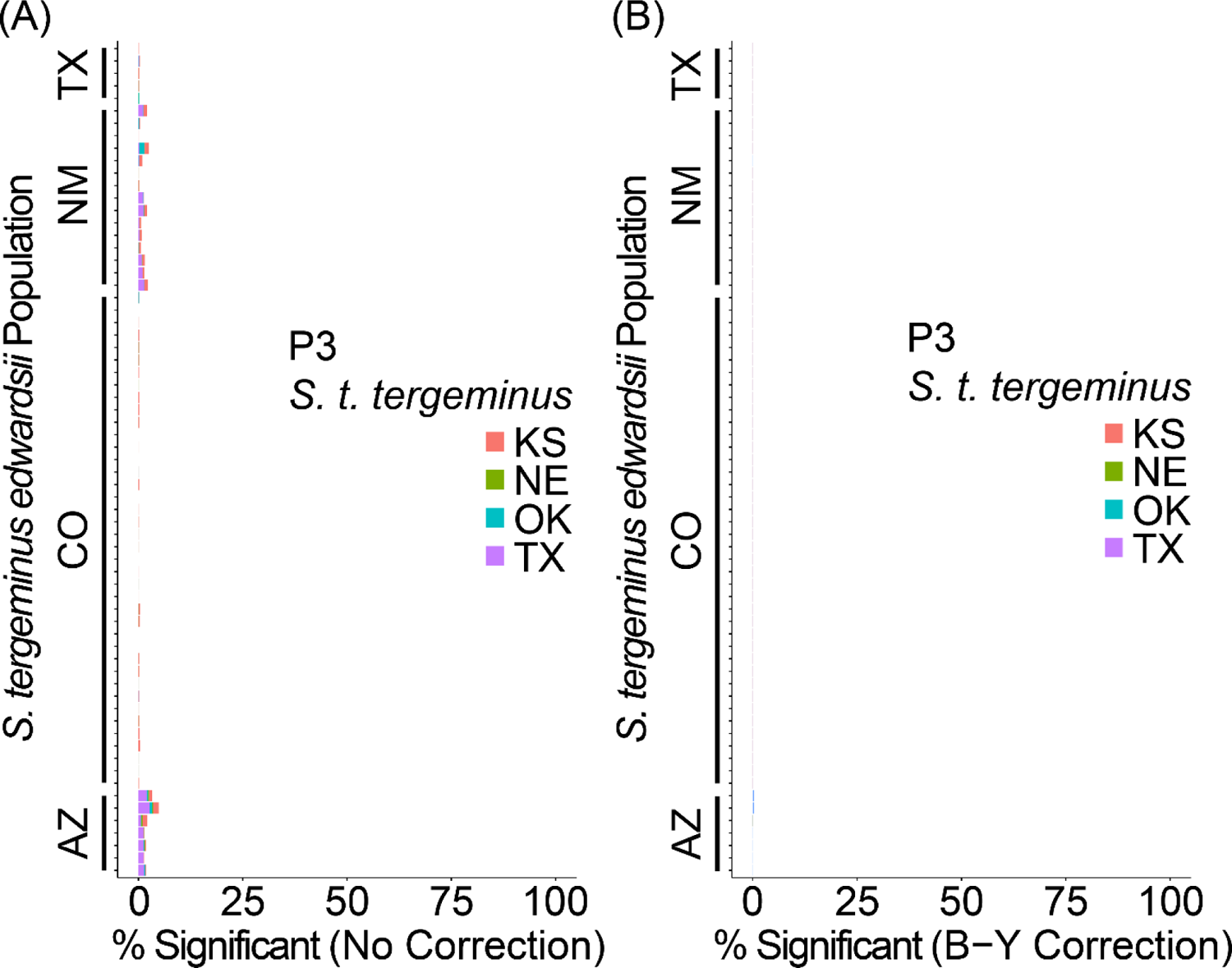
Four-taxon *D*-statistic tests for *Sistrurus tergeminus tergeminus* (P3) and *S. tergeminus edwardsii* (P1 and P2). Stacked bars indicate percent significant *D*-statistic tests per P3 sample locality, with (A) no *P*-value correction for multiple tests and (B) a Benjamini– Yekutieli (B-Y) correction for controlling the false discovery rate. State locality abbreviations include: TX=Texas, NM=New Mexico, CO=Colorado, AZ=Arizona, KS=Kansas, NE=Nebraska, OK=New Oklahoma. The outgroups (P4) included *S. miliarius streckeri* and *S. m. barbouri*.

**Figure S7:**
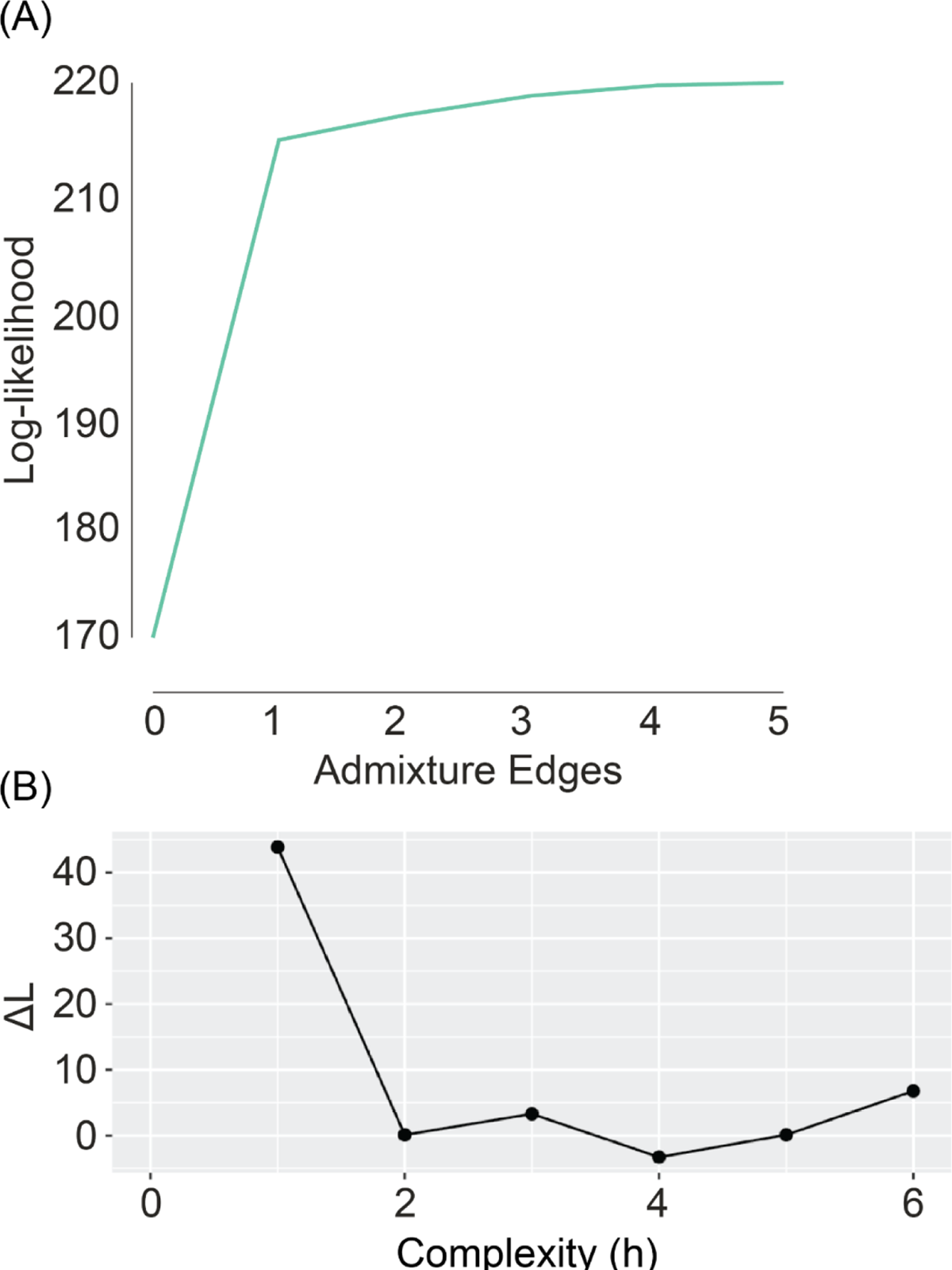
(A) TREEMIX log-likelihoods for 0-5 migration edges. (B) PHYLONETWORKS first-order change in pseudo-likelihood (ΔL) for 0-6 hybrid nodes (h).

## Notes

### Competing Interest Statement

The authors have declared no competing interest.

## References

1. Adams, J., Maslin, M., Thomas, E., 1999. Sudden climate transitions during the Quaternary. Prog. Phys. Geogr. 23, 1–36. https://doi.org/10.1177%2F030913339902300101

2. Alexander, D.H., Lange, K., 2011. Enhancements to the ADMIXTURE algorithm for individual ancestry estimation. BMC Bioinformatics 12, 246. https://doi.org/10.1186/1471-2105-12-246

3. Allendorf, F.W., Leary, R.F., Spruell, P., Wenburg, J.K., 2001. The problems with hybrids: setting conservation guidelines. Trends Ecol. Evol. 16, 613–622. https://doi.org/10.1016/S0169-5347(01)02290-X

4. Anderson, C.D., Gibbs, H.L., Douglas, M.E., Holycross, A.T., 2009. Conservation genetics of the desert massasauga rattlesnake (*Sistrurus catenatus edwardsii*). Copeia 2009, 740–747. https://doi.org/10.1643/CG-08-152

5. Anderson, E., 1949. Introgressive Hybridization. John Wiley and Sons, New York City, NY, USA. https://doi.org/10.5962/bhl.title.4553

6. Andrews, S., 2010. FastQC: a quality control tool for high throughput sequence data. Available online at: http://www.bioinformatics.babraham.ac.uk/projects/fastqc/ [WWW Document].

7. Antonelli, A., 2017. Biogeography: drivers of bioregionalization. Nat. Ecol. Evol. 1, 1–2. https://doi.org/10.1038/s41559-017-0114

8. Axelrod, D.I., 1983. Paleobotanical history of the western deserts, in: Wells, S.G., Haragan, D.R. (Eds.), Origin and Evolution of Deserts. University of New Mexico Press, Albuquerque, NM, USA, pp. 113–129.

9. Axelrod, D.I., 1979. Age and origin of Sonoran Desert vegetation, in: Occasional Papers Vol. 132. California Academy of Sciences, San Francisco, CA, USA, pp. 1–74.

10. Bangs, M.R., Douglas, M.R., Mussmann, S.M., Douglas, M.E., 2018. Unraveling historical introgression and resolving phylogenetic discord within *Catostomus* (Osteichthys: Catostomidae). BMC Evol. Biol. 18, 86. https://doi.org/10.1186/s12862-018-1197-y

11. Bapst, D.W., 2012. paleotree: an R package for paleontological and phylogenetic analyses of evolution. Methods Ecol. Evol. 3, 803–807. https://doi.org/10.1111/j.2041-210x.2012.00223.x

12. Becker, M., Gruenheit, N., Steel, M., Voelckel, C., Deusch, O., Heenan, P.B., McLenachan, P.A., Kardailsky, O., Leigh, J.W., Lockhart, P.J., 2013. Hybridization may facilitate in situ survival of endemic species through periods of climate change. Nat. Clim. Chang. 3, 1039– 1043. https://doi.org/10.1038/nclimate2027

13. Benjamini, Y., Yekutieli, D., 2001. The control of the false discovery rate in multiple testing under xependency. Ann. Stat. 29, 1165–1188. https://doi.org/10.1214/aos/1013699998

14. Bentley Sr, S.J., Blum, M.D., Maloney, J., Pond, L., Paulsell, R., 2016. The Mississippi River source-to-sink system: Perspectives on tectonic, climatic, and anthropogenic influences, Miocene to Anthropocene. Earth-Science Rev. 153, 139–174. https://doi.org/10.1016/j.earscirev.2015.11.001

15. Blomberg, S.P., Garland Jr, T., Ives, A.R., 2003. Testing for phylogenetic signal in comparative data: behavioral traits are more labile. Evolution (N. Y). 57, 717–745. https://doi.org/10.1554/0014-3820(2003)057[0717:tfpsic]2.0.co;2

16. Bohling, J.H., 2016. Strategies to address the conservation threats posed by hybridization and genetic introgression. Biol. Conserv. 203, 321–327. https://doi.org/10.1016/j.biocon.2016.10.011

17. Bonferroni, C.E., 1936. Teoria statistica delle classi e calcolo delle probabilità. Pubbl. del R. Ist. Super. di Sci. Econ. e Commer. di Firenz 8, 3–62.

18. Bylsma, R., Walkup, D.K., Hibbitts, T.J., Ryberg, W.A., Black, A.N., DeWoody, J.A., 2021. Population genetic and genomic analyses of Western Massasauga (*Sistrurus tergeminus* ssp.): implications for subspecies delimitation and conservation. Conserv. Genet. 12, e8599. https://doi.org/10.1007/s10592-021-01420-8

19. CamachoLSanchez, M., VeloLAntón, G., Hanson, J.O., Veríssimo, A., MartínezLSolano, Í., Marques, A., Moritz, C., Carvalho, S.B., 2020. Comparative assessment of rangeLwide patterns of genetic diversity and structure with SNPs and microsatellites: A case study with Iberian amphibians. Ecol. Evol. 10, 10353–10363. https://doi.org/10.22541/au.159542435.52696336

20. Castoe, T.A., Parkinson, C.L., 2006. Bayesian mixed models and the phylogeny of pitvipers (Viperidae: Serpentes). Mol. Phylogenet. Evol. 39, 91–110. https://doi.org/10.1016/j.ympev.2005.12.014

21. Chafin, T.K., Douglas, M.R., Martin, B.T., Douglas, M.E., 2019. Hybridization drives genetic erosion in sympatric desert fishes of western North America. Heredity (Edinb). 123, 759– 773. https://doi.org/10.1038/s41437-019-0259-2

22. Chafin, T.K., Zbinden, Z.D., Douglas, M.R., Martin, B.T., Middaugh, C.R., Gray, M.C., Ballard, J.R., Douglas, M.E., 2021. Spatial population genetics in heavily managed species: Separating patterns of historical translocation from contemporary gene flow in whiteLtailed deer. Evol. Appl. 14, 1673–1689. https://doi.org/10.1111/eva.13233

23. Chan, W.Y., Hoffmann, A.A., van Oppen, M.J.H., 2019. Hybridization as a conservation management tool. Conserv. Lett. 12, e12652. https://doi.org/10.1111/conl.12652

24. Chernomor, O., Von Haeseler, A., Minh, B.Q., 2016. Terrace aware data structure for phylogenomic inference from supermatrices. Syst. Biol. 65, 997–1008.

25. Chiucchi, J.E., Gibbs, H.L., 2010. Similarity of contemporary and historical gene flow among highly fragmented populations of an endangered rattlesnake. Mol. Ecol. 19, 5345–5358.

26. Clark, P.U., MacAyeal, D.R., Andrews, J.T., Bartlein, P.J., 1995. Ice sheets play important role in climate change. Eos, Trans. Am. Geophys. Union 76, 265–270. https://doi.org/10.1029/EO076i027p00265-02

27. Cook, F.R., 1992. After an Ice Age: Zoogeography of the Massasauga within a Canadian Herpetofaunal Perspective, in: Johnson, B., Menzies, V. (Eds.), International Symposium and Workshop on the Conservation of the Eastern Massasauga Rattlesnake. Metropolitan Toronto Zoo, Toronto, CA, pp. 19–25.

28. Cooper, A., Turney, C., Hughen, K.A., Brook, B.W., McDonald, H.G., Bradshaw, C.J.A., 2015. Abrupt warming events drove Late Pleistocene Holarctic megafaunal turnover. Science (80-.). 349, 602–606. https://doi.org/10.1126/science.aac4315

29. Degenhardt, W.G., Painter, C.W., Price, A.H., 2005. Amphibians and Reptiles of New Mexico. University of New Mexico Press, Albuquerque, NM, USA.

30. Donfrancesco, V., Luque-Lora, R., 2021. Managing hybridisation beyond the natural-anthropogenic dichotomy. Conserv. Biol. 2021, 1–3. https://doi.org/10.1111/cobi.13816

31. Douglas, M.E., Douglas, M.R., Schuett, G.W., Porras, L.W., 2009. Climate change and evolution of the New World pitviper genus *Agkistrodon* (Viperidae). J. Biogeogr. 36, 1164–1180. https://doi.org/10.1111/j.1365-2699.2008.02075.x

32. Douglas, M.E., Douglas, M.R., Schuett, G.W., Porras, L.W., 2006. Evolution of rattlesnakes (Viperidae; *Crotalus*) in the warm deserts of western North America shaped by Neogene vicariance and Quaternary climate change. Mol. Ecol. 15, 3353–3374. https://doi.org/10.1111/j.1365-294X.2006.03007.x

33. Draper, D., Laguna, E., Marques, I., 2021. Demystifying negative connotations of hybridization for less biased conservation policies. Front. Ecol. Evol. 9, 268. https://doi.org/10.3389/fevo.2021.637100

34. Briggler, J.T., Johnson, T.R., 2021. The Amphibians and Reptiles of Missouri (3^rd^ Edition). Missouri Department of Conservation, Jefferson City, MO, USA.

35. Durand, E.Y., Patterson, N., Reich, D., Slatkin, M., 2011. Testing for ancient admixture between closely related populations. Mol. Biol. Evol. 28, 2239–2252. https://doi.org/10.1093/molbev/msr048

36. Dussex, N., Taylor, H.R., Irestedt, M., Robertson, B.C., 2018. When genetic and phenotypic data do not agree: the conservation implications of ignoring inconvenient taxonomic evidence. N. Z. J. Ecol. 42, 284–290. https://doi.org/10.20417/nzjecol.42.13

37. Eaton, D.A.R., Overcast, I., 2020. ipyrad: Interactive assembly and analysis of RADseq datasets. Bioinformatics 36, 2592–2594. https://doi.org/10.1093/bioinformatics/btz966

38. Estoup, A., Largiader, C.R., Cornuet, J., Gharbi, K., Presa, P., Guyomard, R., 2000. Juxtaposed microsatellite systems as diagnostic markers for admixture: an empirical evaluation with brown trout (Salmo trutta) as model organism. Mol. Ecol. 9, 1873–1886.

39. Evans, P.D., Gloyd, H.K., 1948. The subspecies of the massasauga, Sistrurus catenatus, in Missouri. Bull. Chicago Acad. Sci. 8, 225–232.

40. Findley, J.S., 1969. Biogeography of southwestern boreal and desert animals, in: Jones Jr., J.K. (Ed.), Contributions in Mammology: A Volume Honering Professor E. Raymond Hall. University of Kansas Press, Lawrence, KS, USA, pp. 113–128.

41. Fitzpatrick, B.M., Ryan, M.E., Johnson, J.R., Corush, J., Carter, E.T., 2015. Hybridization and the species problem in conservation. Curr. Zool. 61, 206–216. https://doi.org/10.1093/czoolo/61.1.206

42. Fontanella, F.M., Feldman, C.R., Siddall, M.E., Burbrink, F.T., 2008. Phylogeography of *Diadophis punctatus*: extensive lineage diversity and repeated patterns of historical demography in a trans-continental snake. Mol. Phylogenet. Evol. 46, 1049–1070. https://doi.org/10.1016/j.ympev.2007.10.017

43. Gerard, D., Gibbs, H.L., Kubatko, L., 2011. Estimating hybridization in the presence of coalescence using phylogenetic intraspecific sampling. BMC Evol. Biol. 11, 291. https://doi.org/10.1186/1471-2148-11-291

44. Gibbs, H.L., Mackessy, S.P., 2009. Functional basis of a molecular adaptation: prey-specific toxic effects of venom from *Sistrurus* rattlesnakes. Toxicon 53, 672–679. https://doi.org/10.1016/j.toxicon.2009.01.034

45. Gibbs, H.L., Murphy, M., Chiucchi, J.E., 2011. Genetic identity of endangered massasauga rattlesnakes (*Sistrurus* sp.) in Missouri. Conserv. Genet. 12, 433–439. https://doi.org/10.1007/s10592-010-0151-3

46. Gibbs, H.L., Sanz, L., Calvete, J.J., 2009. Snake population venomics: proteomics-based analyses of individual variation reveals significant gene regulation effects on venom protein expression in *Sistrurus* rattlesnakes. J. Mol. Evol. 68, 113–125. https://doi.org/10.1007/s00239-008-9186-1

47. Gibbs, H.L., Sanz, L., Sovic, M.G., Calvete, J.J., 2013. Phylogeny-based comparative analysis of venom proteome variation in a clade of rattlesnakes (*Sistrurus* sp.). PLoS One 8, e67220. https://doi.org/10.1371/journal.pone.0067220

48. Green, R.E., Krause, J., Briggs, A.W., Maricic, T., Stenzel, U., Kircher, M., Patterson, N., Li, H., Zhai, W., Fritz, M.H.-Y., 2010. A draft sequence of the Neandertal genome. Science (80-.). 328, 710–722. https://doi.org/10.1126/science.1188021

49. Greene, H.W., 1997. Snakes: The Evolution of Mystery in Nature. University of California Press, Berkeley and Los Angeles, CA, USA.

50. Guiher, T.J., Burbrink, F.T., 2008. Demographic and phylogeographic histories of two venomous North American snakes of the genus *Agkistrodon*. Mol. Phylogenet. Evol. 48, 543–553. https://doi.org/10.1016/j.ympev.2008.04.008

51. Gutenkunst, R.N., Hernandez, R.D., Williamson, S.H., Bustamante, C.D., 2009. Inferring the joint demographic history of multiple populations from multidimensional SNP frequency data. PLoS Genet. 5, e1000695. https://doi.org/10.1371/journal.pgen.1000695

52. Haasl, R.J., Payseur, B.A., 2011. Multi-locus inference of population structure: a comparison between single nucleotide polymorphisms and microsatellites. Heredity (Edinb). 106, 158–171. https://doi.org/10.1038/hdy.2010.21

53. Hammerson, G.A., 2000. Amphibians and Reptiles in Colorado. University Press of Colorado, Louisville, CO, USA.

54. Hein, Steven R., 2015. An Integrative Approach to Species Delimitation in the Massasauga Rattlesnake (Sistrurus Catenatus) with an Emphasis on the Western Massasauga, S. c. tergeminus, and Desert Massasauga, S. c. edwardsii in Texas. Biology Theses. Paper 29. http://hdl.handle.net/10950/313

55. Herman, T.A., Bouzat, J.L., 2016. RangeLwide phylogeography of the fourLtoed salamander: out of Appalachia and into the glacial aftermath. J. Biogeogr. 43, 666–678. https://doi.org/10.1111/jbi.12679

56. Hewitt, G.M., 2001. Speciation, hybrid zones and phylogeography—or seeing genes in space and time. Mol. Ecol. 10, 537–549. https://doi.org/10.1046/j.1365-294x.2001.01202.x

57. Hewitt, G.M., 1999. Post-glacial re-colonization of European biota. Biol. J. Linnaean Soc. 68, 87–112. https://doi.org/10.1111/j.1095-8312.1999.tb01160.x

58. Hewitt, G.M., 1996. Some genetic consequences of ice ages, and their role in divergence and speciation. Biol. J. Linnaean Soc. 58, 247–276. https://doi.org/10.1006/bijl.1996.0035

59. Hewitt, G.M., 1988. Hybrid zones-natural laboratories for evolutionary studies. Trends Ecol. Evol. 3, 158–167.

60. Hoang, D.T., Chernomor, O., von Haeseler, A., Minh, B.Q., Vinh, L.S., 2017. UFBoot2: improving the ultrafast bootstrap approximation. Mol. Biol. Evol. 35, 518–522. https://doi.org/10.1093/molbev/msx281

61. Holman, J.A., 2000. Fossil snakes of North America: origin, evolution, distribution, paleoecology. Indiana University Press, Bloomington, IN, USA.

62. Holmgren, C.A., Penalba, M.C., Rylander, K.A., Betancourt, J.L., 2003. A 16,000 14C yr BP packrat midden series from the USA–Mexico Borderlands. Quat. Res. 60, 319–329. https://doi.org/10.1016/j.yqres.2003.08.001

63. Holycross, A.T., 2002. Conservation biology of two rattlesnakes, Crotalus willardi obscurus and Sistrurus catenatus edwardsii. Arizona State University. https://www.proquest.com/dissertations-theses/conservation-biology-two-rattlesnakes-i-crotalus/docview/304790860/se-2?accountid=8361

64. Holycross, A.T., Mackessy, S.P., 2002. Variation in the diet of *Sistrurus catenatus* (Massasauga), with emphasis on *Sistrurus catenatus edwardsii* (Desert Massasauga). J. Herpetol. 36, 454–464. https://doi.org/10.1670/0022-1511(2002)036[0454:vitdos]2.0.co;2

65. Hunter, K.L., Betancourt, J.L., Riddle, B.R., Van Devender, T.R., Cole, K.L., Spaulding, W.G., 2001. Ploidy race distributions since the Last Glacial Maximum in the North American desert shrub, *Larrea tridentata*. Glob. Ecol. Biogeogr. 10, 521–533. https://doi.org/10.1046/j.1466-822x.2001.00254.x

66. Jackiw, R.N., Mandil, G., Hager, H.A., 2015. A framework to guide the conservation of species hybrids based on ethical and ecological considerations. Conserv. Biol. 29, 1040–1051. https://doi.org/10.1111/cobi.12526

67. Jones, M.R., Mills, L.S., Jensen, J.D., Good, J.M., 2020. The origin and spread of locally adaptive seasonal camouflage in snowshoe hares. Am. Nat. 196, 316–332.

68. Jouganous, J., Long, W., Ragsdale, A.P., Gravel, S., 2017. Inferring the joint demographic history of multiple populations: beyond the diffusion approximation. Genetics 206, 1549– 1567. https://doi.org/10.1101/103275

69. Kalyaanamoorthy, S., Minh, B.Q., Wong, T.K.F., von Haeseler, A., Jermiin, L.S., 2017. ModelFinder: Fast model selection for accurate phylogenetic estimates. Nat. Methods 14, 587–589. https://doi.org/10.1038/nmeth.4285

70. Kareiva, P., Marvier, M., 2012. What is conservation science? Bioscience 62, 962–969.

71. Kembel, S.W., Cowan, P.D., Helmus, M.R., Cornwell, W.K., Morlon, H., Ackerly, D.D., Blomberg, S.P., Webb, C.O., 2010. Picante: R tools for integrating phylogenies and ecology. Bioinformatics 26, 1463–1464. https://doi.org/10.1093/bioinformatics/btq166

72. Kubatko, L.S., Gibbs, H.L., Bloomquist, E.W., 2011. Inferring species-level phylogenies and taxonomic distinctiveness using multilocus data in *Sistrurus* rattlesnakes. Syst. Biol. 60, 393–409. https://doi.org/10.1093/sysbio/syr011

73. Larget, B.R., Kotha, S.K., Dewey, C.N., Ané, C., 2010. BUCKy: Gene tree/species tree reconciliation with Bayesian concordance analysis. Bioinformatics 26, 2910–2911. https://doi.org/10.1093/bioinformatics/btq539

74. Lee, K.M., Kivelä, S.M., Ivanov, V., Hausmann, A., Kaila, L., Wahlberg, N., Mutanen, M., 2018. Information dropout patterns in RAD phylogenomics and a comparison with multilocus Sanger data in a species-rich moth genus. Syst. Biol. 67, 925–939. https://doi.org/10.1093/sysbio/syy029

75. Linck, E.B., Battey, C.J., 2019. Minor allele frequency thresholds strongly affect population structure inference with genomic datasets. Mol. Ecol. Resour. 19, 639–647. https://doi.org/10.1111/1755-0998.12995

76. Lowe, C.H., Schwalbe, C.R., Johnson, T.B., 1986. The Venomous Reptiles of Arizona. Arizona Game and Fish Department, Phoenix, AZ, USA.

77. MacKay, W.P., Elias, S.A., 1992. Late Quaternary ant fossils from packrat middens (Hymenoptera: Formicidae): implications for climatic change in the Chihuahuan Desert. Psyche (Stuttg). 99, 169–184. https://doi.org/10.1155/1992/82014

78. Mackessy, S., 2005. Desert Massasauga Rattlesnake (Sistrurus catenatus edwardsii): a technical conservation assessment. USDA Forest Service, Rocky Mountain Region. Available http://www.fs.fed.us/r2/projects/scp/assessments/massasauga.pdf. (Accessed January 10, 2021). [WWW Document]. URL http://www.fs.fed.us/r2/projects/scp/assessments/massasauga.pdf (accessed 1.10.21).

79. Martin, B.T., Douglas, M.R., Chafin, T.K., Placyk, J.S., Birkhead, R.D., Phillips, C.A., Douglas, M.E., 2020. Contrasting signatures of introgression in North American box turtle (*Terrapene* spp.) contact zones. Mol. Ecol. 29, 4186–4202. https://doi.org/10.1111/mec.15622

80. McCartney-Melstad, E., Gidiş, M., Shaffer, H.B., 2019. An empirical pipeline for choosing the optimal clustering threshold in RADseq studies. Mol. Ecol. Resour. 19, 1195–1204. https://doi.org/10.1111/1755-0998.13029

81. McCluskey, E.M., Bender, D., 2015. Genetic structure of western massasauga rattlesnakes (*Sistrurus catenatus tergeminus*). J. Herpetol. 49, 343–348. https://doi.org/10.1670/14-016

82. Minh, B.Q., Schmidt, H.A., Chernomor, O., Schrempf, D., Woodhams, M.D., Von Haeseler, A., Lanfear, R., 2020. IQ-TREE 2: New models and efficient methods for phylogenetic inference in the genomic era. Mol. Biol. Evol. 37, 1530–1534. https://doi.org/10.1093/molbev/msaa015

83. Minton, S.A., 1983. *Sistrurus catenatus* (Rafinesque). Cat. Am. Amphib. Reptil. (CAAR), Soc. Study Amphib. Reptil. 332.1–332.2.

84. Morafka, D.J., 1977. A biogeographical analysis of the Chihuahuan desert through its herpetofauna. Biogeographica 9, 1–313. https://doi.org/10.1007/978-94-010-1318-5

85. Morin, P.A., Luikart, G., Wayne, R.K., 2004. SNPs in ecology, evolution and conservation. Trends Ecol. Evol. Evol. 19, 208–216. https://doi.org/10.1016/j.tree.2004.01.009

86. Murphy, R.W., Fu, J., Lathrop, A., Feltham, J. V, Kovac, V., 2002. Phylogeny of the rattlesnakes (Crotalus and Sistrurus) inferred from sequences of five mitochondrial DNA genes, in: Schuett, G.W., Höggren, M., Douglas, M.E., Greene, H.W. (Eds.), Biology of the Vipers. Eagle Mountain Press LC, Salt Lake City, UT, USA, pp. 69–92.

87. Mussmann, S.M., Douglas, M.R., Bangs, M.R., Douglas, M.E., 2020a. Comp-D: a program for comprehensive computation of D-statistics and population summaries of reticulated evolution. Conserv. Genet. Resour. 12, 263–267. https://doi.org/10.1007/s12686-019-01087-x

88. Mussmann, S.M., Douglas, M.R., Chafin, T.K., Douglas, M.E., 2020b. AdmixPipe: population analyses in Admixture for non-model organisms. BMC Bioinformatics 21, 1–9.

89. National Research Council, 2002. Abrupt climate change: inevitable surprises. National Academy Press, Washington, D.C., USA. https://doi.org/10.17226/10136.

90. Neri-Castro, E., Strickland, J.L., Carbajal-Márquez, R.A., Zuñiga, J., Ponce-López, R., Olvera, F., Alagón, A., 2022. Characterization of the venom and external morphology of a natural hybrid between *Crotalus atrox* and *Crotalus mictlantecuhtli*. Toxicon 207, 43–47. https://doi.org/10.1016/j.toxicon.2022.01.003

91. Noskova, E., Ulyantsev, V., Koepfli, K.-P., O’Brien, S.J., Dobrynin, P., 2020. GADMA: Genetic algorithm for inferring demographic history of multiple populations from allele frequency spectrum data. Gigascience 9, giaa005. https://doi.org/10.1093/gigascience/giaa005

92. Ochoa, A., Broe, M., Moriarty Lemmon, E., Lemmon, A.R., Rokyta, D.R., Gibbs, H.L., 2020. Drift, selection and adaptive variation in small populations of a threatened rattlesnake. Mol. Ecol. 29, 2612–2625. https://doi.org/10.1111/mec.15517

93. Ottenburghs, J., 2021. The genic view of hybridization in the Anthropocene. Evol. Appl. 14, 2342–2360. https://doi.org/10.1111/eva.13223

94. Pacheco-Sierra, G., Vázquez-Domínguez, E., Pérez-Alquicira, J., Suárez-Atilano, M., Domínguez-Laso, J., 2018. Ancestral hybridization yields evolutionary distinct hybrids lineages and species boundaries in crocodiles, posing unique conservation conundrums. Front. Ecol. Evol. 6, 138. https://doi.org/10.3389/fevo.2018.00138

95. Parkinson C.L., Chippindale, P., Campbell, J., 2002. Multigene analyses of pitviper phylogeny with comments on their biogeographical history. In: Schuett, G.W., Hoggren, M., Douglas, M.E., Greene, H.W. (Ed.). Biology of the vipers. Eagle Mountain (Utah): Eagle Mountain Publishing, LC. p. 93–110.

96. Parmley, D., Holman, J.A., 2007. Earliest fossil record of a pigmy rattlesnake (Viperidae: *Sistrurus* Garman). J. Herpetol. 41, 141–144. https://doi.org/10.1670/0022-1511(2007)41[141:EFROAP]2.0.CO;2

97. Payseur, B.A., 2010. Using differential introgression in hybrid zones to identify genomic regions involved in speciation. Mol. Ecol. Resour. 10, 806–820. https://doi.org/10.1111/j.1755-0998.2010.02883.x

98. Pendall, E., Betancourt, J.L., Leavitt, S.W., 1999. Paleoclimatic significance of δD and δ13C values in piñon pine needles from packrat middens spanning the last 40,000 years. Palaeogeogr. Palaeoclimatol. Palaeoecol. 147, 53–72. https://doi.org/10.1016/S0031-0182(98)00152-7

99. Pickrell, J.K., Pritchard, J.K., 2012. Inference of population splits and mixtures from genome-wide allele frequency data. PLoS Genet. 8, e1002967. https://doi.org/10.1038/npre.2012.6956.1

100. QGIS Development Team 2020 QGIS Geographic Information System. Open Source Geospatial Foundation Project. http://qgis.osgeo.org [WWW Document], n.d. URL http://qgis.org

101. R Development Core Team, 2018. R: A language and environment for statistical computing. https://cran.r-project.org/.

102. Rašić, G., Filipović, I., Weeks, A.R., Hoffmann, A.A., 2014. Genome-wide SNPs lead to strong signals of geographic structure and relatedness patterns in the major arbovirus vector, *Aedes aegypti*. BMC Genomics 15, 275. https://doi.org/10.1186/1471-2164-15-275

103. Remington, C.L., 1968. Suture-zones of hybrid interaction between recently joined biotas, in: Dobzhansky, T., Hecht, M.K., Steere, W.C. (Ed.), Evolutionary Biology. Springer, Boston, MA, USA, pp. 321–428. https://doi.org/10.1007/978-1-4684-8094-8_8

104. Revell, L.J., 2012. phytools: an R package for phylogenetic comparative biology (and other things). Methods Ecol. Evol. 3, 217–223. https://doi.org/10.1111/j.2041-210X.2011.00169.x

105. Rhymer, J.M., Simberloff, D., 1996. Extinction by hybridization and introgression. Annu. Rev. Ecol. Syst. 27, 83–109. https://doi.org/10.1146/annurev.ecolsys.27.1.83

106. Ronquist, F., Teslenko, M., van der Mark, P., Ayres, D.L., Darling, A., Höhna, S., Larget, B., Liu, L., Suchard, M.A., Huelsenbeck, J.P., 2012. MrBayes 3.2: efficient Bayesian phylogenetic inference and model choice across a large model space. Syst. Biol. 61, 539–542. https://doi.org/10.1093/sysbio/sys029

107. Rozas, J., Ferrer-Mata, A., Sánchez-DelBarrio, J.C., Guirao-Rico, S., Librado, P., Ramos-Onsins, S.E., Sánchez-Gracia, A., 2017. DnaSP 6: DNA sequence polymorphism analysis of large data sets. Mol. Biol. Evol. 34, 3299–3302. https://doi.org/10.1093/molbev/msx248

108. Ruane, S., Bryson Jr, R.W., Pyron, R.A., Burbrink, F.T., 2014. Coalescent species delimitation in milksnakes (genus *Lampropeltis*) and impacts on phylogenetic comparative analyses. Syst. Biol. 63, 231–250. https://doi.org/10.1093/sysbio/syt099

109. Ryberg, W.A., Harvey, J.A., Blick, A., Hibbitts, T.J., Voelker, G., 2015. Genetic structure is inconsistent with subspecies designations in the western massasauga *Sistrurus tergeminus*. J. Fish Wildl. Manag. 6, 350–359. https://doi.org/10.3996/122014-JFWM-093

110. Sanz, L., Gibbs, H.L., Mackessy, S.P., Calvete, J.J., 2006. Venom proteomes of closely related *Sistrurus* rattlesnakes with divergent diets. J. Proteome Res. 5, 2098–2112. https://doi.org/10.1021/pr0602500

111. Sartor, C.C., Cushman, S.A., Wan, H.Y., Kretschmer, R., Pereira, J.A., Bou, N., Cosse, M., González, S., Eizirik, E., de Freitas, T.R.O., 2021. The role of the environment in the spatial dynamics of an extensive hybrid zone between two neotropical cats. J. Evol. Biol. 34, 614– 627. https://doi.org/10.1111/jeb.13761

112. Savage, J.M., 1960. Evolution of a peninsular herpetofauna. Syst. Zool. 9, 184–212. https://doi.org/10.2307/2411967

113. Seigel, R.A., 1986. Ecology and conservation of an endangered rattlesnake, *Sistrurus catenatus*, in Missouri, USA. Biol. Conserv. 35, 333–346. https://doi.org/10.1016/0006-3207(86)90093-5

114. Snir, S., Rao, S., 2012. Quartet MaxCut: A fast algorithm for amalgamating quartet trees. Mol. Phylogenet. Evol. 61, 1–8. https://doi.org/10.1016/j.ympev.2011.06.021

115. Solís-Lemus, C., Ané, C., 2016. Inferring phylogenetic networks with Maximum Pseudolikelihood under incomplete lineage sorting. PLoS Genet. 12, 1–21. https://doi.org/10.1371/journal.pgen.1005896

116. Solís-Lemus, C., Bastide, P., Ané, C., 2017. PhyloNetworks: A package for phylogenetic networks. Mol. Biol. Evol. 34, 3292–3298. https://doi.org/10.1093/molbev/msx235

117. Soulé, M., 2013. The “New Conservation.” Conserv. Biol. 28, 895–897. https://doi.org/10.5822/978-1-61091-559-5_7

118. Sovic, M., Fries, A., Martin, S.A., Lisle Gibbs, H., 2019. Genetic signatures of small effective population sizes and demographic declines in an endangered rattlesnake, *Sistrurus catenatus*. Evol. Appl. 12, 664–678. https://doi.org/10.1111/eva.12731

119. Sovic, M.G., Fries, A.C., Gibbs, H.L., 2016. Origin of a cryptic lineage in a threatened reptile through isolation and historical hybridization. Heredity (Edinb). 117, 358–366. https://doi.org/10.1038/hdy.2016.56

120. Stebbins, R.C., 2003. A Field Guide to Western Reptiles and Amphibians, 3rd ed. Houghton Mifflin Harcourt, Boston, MA, USA.

121. Stenz, N.W.M., Larget, B., Baum, D.A., Ané, C., 2015. Exploring tree-like and non-tree-like patterns using genome sequences: An example using the inbreeding plant species *Arabidopsis thaliana* (L.) Heynh. Syst. Biol. 64, 809–823. https://doi.org/10.1093/sysbio/syv039

122. Sunde, J., Yıldırım, Y., Tibblin, P., Forsman, A., 2020. Comparing the performance of microsatellites and RADseq in population genetic studies: Analysis of data for pike (*Esox lucius*) and a synthesis of previous studies. Front. Genet. 11, 218. https://doi.org/10.3389/fgene.2020.00218

123. Swenson, N.G., Howard, D.J., 2005. Clustering of contact zones, hybrid zones, and phylogeographic breaks in North America. Am. Nat. 166, 581–591. https://doi.org/10.1086/491688

124. Szatmári, L., Cserkész, T., Laczkó, L., Lanszki, J., Pertoldi, C., Abramov, A. V, Elmeros, M., Ottlecz, B., Hegyeli, Z., Sramkó, G., 2021. A comparison of microsatellites and genomeLwide SNPs for the detection of admixture brings the first molecular evidence for hybridization between *Mustela eversmanii* and *M. putorius* (Mustelidae, Carnivora). Evol. Appl. 14, 2286–2304. https://doi.org/10.1111/eva.13291

125. Szymanski, J., 1998. Status assessment for the eastern massasauga (*Sistrurus c. catenatus*). U.S. Fish and Wildlife Service. Fort Snelling, MN, USA.

126. Szymanski, J., Pollack, C., Ragan, L., Redmer, M., Clemency, L., Voorhies, K., Jaka, J., 2016. Species status assessment for the eastern massasauga rattlesnake (*Sistrurus catenatus*). U.S. Fish and Wildlife Service. Fort Snelling, MN, USA.

127. Taylor, S.A., Larson, E.L., 2019. Insights from genomes into the evolutionary importance and prevalence of hybridization in nature. Nat. Ecol. Evol. 3, 170–177.

128. Taylor, S.A., Larson, E.L., Harrison, R.G., 2015. Hybrid zones: windows on climate change. Trends Ecol. Evol. 30, 398–406.

129. To, T.-H., Jung, M., Lycett, S., Gascuel, O., 2016. Fast dating using least-squares criteria and algorithms. Syst. Biol. 65, 82–97.

130. Van Devender, T.R., 1977. Holocene woodlands in the southwestern deserts. Science (80-.). 198, 189–192. https://doi.org/10.1126/science.198.4313.189

131. Vendrami, D.L.J., Telesca, L., Weigand, H., Weiss, M., Fawcett, K., Lehman, K., Clark, M.S., Leese, F., McMinn, C., Moore, H., 2017. RAD sequencing resolves fine-scale population structure in a benthic invertebrate: implications for understanding phenotypic plasticity. R. Soc. Open Sci. 4, 160548.

132. Von Holdt, B.M., Brzeski, K.E., Wilcove, D.S., Rutledge, L.Y., 2018. Redefining the role of admixture and genomics in species conservation. Conserv. Lett. 11, e12371.

133. Walkup, D.K., Lawing, A.M., Hibbitts, T.J., Ryberg, W.A., 2022. Biogeographic consequences of shifting climate for the Western Massasauga (*Sistrurus tergeminus*). Ecol. Evol. 12, e8599.

134. Wastell, A.R., Mackessy, S.P., 2016. Desert Massasauga Rattlesnakes (*Sistrurus catenatus edwardsii*) in southeastern Colorado: life history, reproduction, and communal hibernation. J. Herpetol. 50, 594–603.

135. Wastell, A.R., Mackessy, S.P., 2011. Spatial ecology and factors influencing movement patterns of Desert Massasauga rattlesnakes (*Sistrurus catenatus edwardsii*) in southeastern Colorado. Copeia 2011, 29–37.

136. Werler, J.E., Dixon, J.R., 2010. Texas Snakes: Identification, Distribution, and Natural History. University of Texas Press, Austin, TX, USA.

137. Wilson, J.S., Pitts, J.P., 2010. Illuminating the lack of consensus among descriptions of earth history data in the North American deserts: a resource for biologists. Prog. Phys. Geogr. 34, 419–441.

138. Wooten, J.A., Gibbs, H.L., 2012. Niche divergence and lineage diversification among closely related *Sistrurus* rattlesnakes. J. Evol. Biol. 25, 317–328.

139. Yu, G., Smith, D.K., Zhu, H., Guan, Y., Lam, T.T., 2017. ggtree: an R package for visualization and annotation of phylogenetic trees with their covariates and other associated data. Methods Ecol. Evol. 8, 28–36.

140. Zecherle, L.J., Nichols, H.J., BarLDavid, S., Brown, R.P., Hipperson, H., Horsburgh, G.J., Templeton, A.R., 2021. Subspecies hybridization as a potential conservation tool in species reintroductions. Evol. Appl. 14, 1216–1224.

141. Zimmerman, S.J., Aldridge, C.L., Oyler-McCance, S.J., 2020. An empirical comparison of population genetic analyses using microsatellite and SNP data for a species of conservation concern. BMC Genomics 21, 1–16.

